# An isogenic human iPSC model unravels neurodevelopmental abnormalities in SMA

**DOI:** 10.1101/2023.01.02.522499

**Authors:** Tobias Grass, Ines Rosignol, Joshua Thomas, Felix Buchner, Zeynep Dokuzluoglu, Anna Dalinskaya, Jutta Becker, Brunhilde Wirth, Natalia Rodriguez-Muela

## Abstract

Whether neurodevelopmental defects underlie the selective neuronal death that characterizes neurodegenerative diseases is becoming an intriguing question. To address it, we focused on the motor neuron (MN) disease Spinal Muscular Atrophy (SMA), caused by reduced levels of the ubiquitous protein SMN. Taking advantage of the first isogenic human induced pluripotent stem cell-derived SMA model that we have generated and a spinal cord organoid system, here we report that the relative and temporal expression of early neural progenitor and MN markers is altered in SMA. Furthermore, the corrected isogenic controls only partially reverse these abnormalities. These findings raise the relevant clinical implication that SMN-increasing treatments might not fully amend SMA pathological phenotypes. The approach we have taken demonstrates that the discovery of new disease mechanisms is greatly improved by using human isogenic models. Moreover, our study implies that SMA has a developmental component that might trigger the MN degeneration.

## Introduction

Accumulating evidence suggests that potentially all neurodegenerative diseases (NDDs) have a developmental component that could be essential for postmitotic neurons to manifest disease hallmarks. Yet, even in the cases where a known mutated or deleted gene is the cause of the disease (i.e. familial forms of Amyotrophic Lateral Sclerosis, Alzheimer’s and Parkinson’s diseases or the fully genetic Huntington’s Disease) it remains a mystery why the pathology only manifests years after birth, often after decades. Probably for this reason, these diseases have not been studied from a developmental angle. However, recent, yet scarce, evidence is revealing unanticipated neurodevelopmental alterations that could swift the way these NDDs are understood.

Spinal Muscular Atrophy (SMA) is an autosomal, recessively inherited, neuromuscular disease where spinal motoneurons (MNs) primarily degenerate leading to muscle wasting and premature death in the most severe cases. It is caused by mutations or full deletion of the *SMN1* gene, which codes for the survival of motoneuron (SMN) protein (Lefebvre et al., 1995; Lefebvre et al., 1997). Only in humans, a second paralogous gene exists, *SMN2*, which partially compensates for the *SMN1* loss allowing SMA patient survival. *SMN2* is almost identical to *SMN1* but, importantly, carries a translationally silent point mutation in exon 7 that disrupts its splicing (Lorson et al., 1999). As a result, only a small fraction of the *SMN2* transcripts produce full-length SMN protein, while the majority of *SMN2* products lack exon 7 and generate a truncated, unstable protein (Cho and Dreyfuss, 2010; Lorson et al., 1998). *SMN2* copy number varies between individuals, and therefore the amount of *SMN2*-derived full-length SMN protein does too, which accounts, at least partially, for a wide spectrum of SMA disease severities (Harada et al., 2002; Feldkotter et al., 2002).

SMN is an ubiquitous and key protein for the assembly of spliceosomal small nuclear ribonucleoproteins (snRNPs) that mediate pre-mRNA splicing (Gubitz et al., 2004). Complete lack of SMN results in cell death (Cifuentes-Diaz et al., 2001; Vitte et al., 2004) and *SMN* knockout mice die at the morula stage (Schrank et al., 1997). The nuclear chaperone role of SMN was the first described and is the best molecularly characterized SMN function to date. However, SMN has been involved in numerous other intracellular pathways, such as axonal transport of mRNAs and RNPs, ribosomal dynamics and translation (Lauria et al., 2020), mitochondrial trafficking (James et al., 2021; Zilio et al., 2022), endosomal and membrane recycling pathways (Dimitriadi et al., 2016; Riessland et al., 2017; Hosseinibarkooie et al., 2016) and autophagy (Rodriguez-Muela et al., 2018; Sansa et al., 2021). SMN binds to a large variety of proteins engaged in multiple cellular mechanisms; it is therefore expected that a better understanding of the protein’s biology will uncover additional SMN functions. It is also possible that anomalies in any of those pathways upon SMN deficiency derive into a neural progenitor defect that gives rise to neurons poised to degenerate later in life. While the less severe SMA types III and IV display clinical manifestations fitting with a classical NDD, the most severe types 0-I present features consistent with a developmental disease (Wirth, 2021). Examples of these are developmentally immature motor axons in SMA fetuses and SMA mouse model embryos, which fail to reach a proper radial growth and myelination during embryogenesis (Kong et al., 2021). Further, depleting SMN in Olig2+ MN progenitors causes a SMA-like phenotype in mice (Park et al., 2010). Additionally, reducing or increasing SMN levels selectively in neuroblasts in a defined time-frame manner, prior to maturation and establishment of neuromuscular junctions, impaired or restored locomotor function in *Drosophila*, respectively (Grice and Liu, 2022). It is also well established that SMN protein levels are higher and more strongly required during the early stages of prenatal development than postnatally (Le et al., 2011; Foust et al., 2010; Hua et al., 2010; Hua et al., 2011; Porensky et al., 2012; Bogdanik et al., 2015; Lutz et al., 2011), and that accordingly, early therapeutic interventions are needed in order to achieve better clinical outcomes (Zhou et al., 2015; Dangouloff and Servais, 2019; Martinez-Hernandez et al., 2013).

The use of human derived *in vitro* models to study human neurodevelopment and disease has been greatly exploited in the recent years (Uzquiano and Arlotta, 2022; Kelley and Pasca, 2022). Cerebral organoids have synergized with model organism research to deepen our understanding of human physiology and pathology and, importantly, provide great translatability to clinical trials emerging as powerful tools to discover and test new therapeutic candidates. While they are being extensively used to model and study numerous NDDs (Del Dosso et al., 2020), their application in SMA research is just beginning (Hor et al., 2018). Nonetheless, an important limitation that human-based research still faces is that very few studies are performed in an isogenic context. Healthy control and disease hiPSCs are generated from somatic cells collected from different donors having, therefore, different genetic backgrounds. How those genetic differences impact disease phenotypes is unknown, however it has certainly affected reproducibility of hiPSC-based experiments and, more importantly, it could lead to inaccurate or even misleading findings (Volpato and Webber, 2020).

The first aim of this study was to generate an isogenic SMA human model, composed of a cohort of three disease lines generated from patients affected by different severities, and two lines derived from each of them engineered to differ at only the silent mutation in one *SMN2* locus responsible for the exon 7 skipping (Lorson et al., 1999). To achieve this a knock-in, CRISPR/Cas9 and vector-based genome editing approach was successfully pursued, by which the green fluorescent protein Clover was also introduced downstream of the corrected *SMN2* exon 7. This constituted an additional advantage to the isogeneity as it allowed the visualization and tracking of SMN protein in live-imaging experiments. To our knowledge, this is the first study to produce a battery of CRISPR/Cas9-corrected human SMA lines and the first one where SMN protein has been tagged endogenously with a fluorescent reporter in hiPSCs. Using this advanced model, we hypothesized and unraveled that SMN deficiency alters the development of neural progenitors, which could sensitize specific MN populations to succumb later as the disease progresses. Taking advantage of a spinal organoid (SCO) model that we have newly developed, this isogenic model has enabled us to identify numerous abnormalities at the stem cell, neural progenitor and MN level that, interestingly, were only partially amended in the corrected isogenic clones.

Our study has two major implications: first, only by comparing disease and isogenic corrected hiPSC-derived SCOs certain disease-specific mechanisms that indicate a developmental aberration in the SMA spinal cord could be identified, which is essential not only for understanding the pathology but also to design new therapies. Second, the developmental defects observed in the SMA SCOs at the level of neural progenitors and postmitotic MNs were not fully reversed in the corrected clones. These findings suggest that restoring SMN protein levels in SMA patients – regardless of how early in development– might not be enough to fully correct disease phenotypes. Further studies will be needed to fully understand the nature and extent of the neurodevelopmental abnormalities occurring upon SMN deficiency and to determine whether these defects in the spinal cord development underlie the selective MN death that characterizes SMA pathology.

## Results

### Generation of isogenic control hiPSCs from various severities of SMA hiPSC lines

We have used three previously described human SMA iPSC lines (Ng et al., 2015; Rodriguez-Muela et al., 2017; Rodriguez-Muela et al., 2018) and a CRISPR/Cas9-mediated genome editing approach to generate three isogenic hiPSCs trios, where the T to C mutation in position 6 of exon 7 in at least one of the *SMN2* copies in each SMA line has been corrected. One SMA type I (named 38D-I for short), one type II (51N-II) and one type III (39C-III) hiPSC lines were edited following a two-vector based, knock-in targeting-approach (Ahfeldt et al., 2020) (Figure 1A-B). Two donor DNA fragments were used, the first consisting of 440 bp homologous to the last 372 bp of intron 6-exon 7, including the corrected exon 7 mutation and excluding exon 7 stop codon, and a second consisting of 440 bp homologous to the 3’ region downstream of exon 7 and containing the coding sequence of the reporter *Clover* -an enhanced fluorescent version of eGFP (Lam et al., 2012)-(Figure S1A). MLPA (Multiplex-Ligation-dependent Probe Amplification) from genomic DNA of the converted hiPSC clones failed to identify the converted *SMN1* due to inability of the probes to bind after *SMN2* conversion into *SMN1:Clover*. Instead, Sanger sequencing confirmed the correct integration of the donor DNA and therefore the conversion of at least one *SMN2* gene into a *SMN1* (Figure 1C, Figure S1D-E). PCR results also showed that at least one *SMN2* copy remained untargeted in each of the lines (Figure S1B-C). Two clones per line were selected for subsequent studies. All analyzed clones were karyotypically normal (data not shown) and expressed normal levels of standard pluripotency markers (OCT4, SOX2, NANOG, TRA160) (Figure S1F-G). We have previously shown that SMN protein levels, measured by western blot from spinal MNs lysates differentiated from these 3 SMA hiPSCs, is ∼10% for the type I compared to healthy control lines and ∼40% for the types II and III (Rodriguez-Muela et al., 2017). Western blot analysis of the two clones and their respective parental line confirmed the presence of only untagged SMN protein in the parental lines and high levels of SMN:Clover in the corrected clones (Figure 1D-F). A healthy control line, BJ siPSD (named “BJ” for short) (Ng et al., 2015; Rodriguez-Muela et al., 2017; Rodriguez-Muela et al., 2018), was used as an additional control for some experiments, and was genetically edited with a similar approach (Figure 1B).

**Figure 1.**
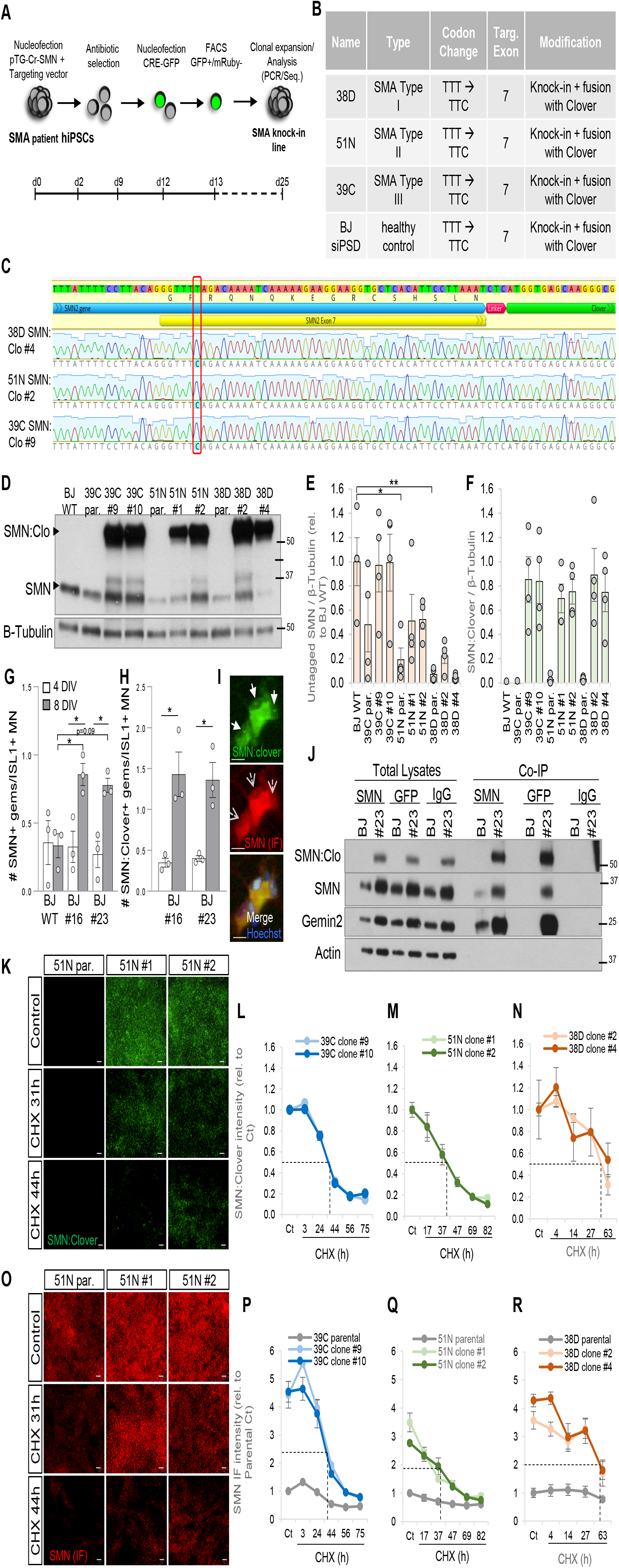
Generation of isogenic control hiPSCs from various severities of SMA human iPSC lines. (**A**) Knock-in CRISPR/Cas9-mediated mutagenesis workflow to correct the C to T mutation in SMA hiPSC lines. (**B**) Isogenic corrected hiPSC lines generated and used for the study. Codon change is specific for the changes made to the *SMN2* gene upon targeting with the CRISPR/Cas9 constructs. (**C**) Sanger-sequencing results from the successfully targeted SMA lines aligned to the exon 7 of *SMN2* gene. Chromatograms from one clone of each successfully targeted SMA line highlighting the corrected C in position 6 of exon 7 are shown (type I “38D-SMN:Clo #4”; type II “51N-SMN:Clo #2” and type III “39C-SMN:Clo #9”). (**D**) Representative western blot from hiPSC lysates from the 3 isogenic SMA trios showing SMN:Clover and SMN protein levels and untagged SMN (**E**) and SMN:Clover (**F**) quantifications. SCO (*p<0.05, **p<0.01 One-way ANOVA followed by Tukey’s multiple comparison test, n=4). (**G**) Quantification of the number of nuclear gems endogenously identified by SMN:Clover or immunostained using an anti-SMN antibody (**H**) in MN cultures generated from the healthy control BJ WT line and two SMN:Clover edited clones, 4 and 8 days after being plated (“BJ WT #16” and “BJ WT #23”). SCO (*p<0.05 One-way ANOVA followed by Tukey’s multiple comparison test, n=3). Representative images indicating SMN:Clover+ (filled arrows) and SMN immunostained nuclear gems (empty arrows) are shown in (**I**). Scale bar, 10 µm. (**J**) Representative immunoprecipitation from BJ WT and BJ SMN:Clover #23 hiPSC lysates. Anti-SMN or anti-GF nanobody-coated beads were used to pull down SMN or Clover (anti-IgG antibody was used as control). SMN:Clover, SMN and Gemin2 proteins are shown. (**K**) Representative images of endogenous SMN:Clover on the isogenic SMA type II hiPSCs trio (51N parental line and corrected clones #1 and #2) treated with 0.3µg/ml cycloheximide (CHX) for the indicated times versus control (DMSO). Scale bar, 50 µm. Quantification of the SMN:Clover fluorescence intensity (proxy for protein levels) in hiPSCs treated with CHX for the indicated times (in hours) related to the DMSO-treated cells (Ct) to determine SMN:Clover half-life in the SMA 39C-III (**L**), 51N-II (**M**) and 38D-I (**N**) hiPSCs. (**O**) Representative images of immunostained SMN total levels on the isogenic SMA type II hiPSCs trio treated with CHX for the indicated times versus control. Scale bar, 50 µm. Quantification of total SMN protein levels related to the DMSO-treated cells to determine total (untagged and SMN:Clover) half-life in the SMA 39C-III (**P**), 51N-II (**Q**) and 38D-I (**R**) hiPSCs.

### The C-terminus Clover reporter does not affect the main properties of SMN protein

To rule out whether in-frame addition of the Clover reporter to the full-length SMN C-terminus altered the biology of the protein, numerous quality control assays were performed. SMN is localized both in the cytoplasm and nuclear foci termed gems, which are frequently adjacent to or overlapping with Cajal bodies (Cauchi, 2010), where it plays a key chaperone role in the biogenesis of snRNPs. Gems are multiprotein nuclear structures composed of SMN proteins, gemins2-7 and snRNPs involved in the transcription and processing of nuclear RNAs (Gubitz et al., 2004). First, we explored whether the addition of the tag would influence SMN recruitment to nuclear gems or their formation. The BJ parental line and two BJ edited hiPSC clones were differentiated into spinal MNs following a widely used embryoid-body (EB)-based differentiation protocol (Rodriguez-Muela et al., 2017; Wichterle et al., 2002) and the resulting MN cultures were fixed and immunostained against SMN 4 and 8 days after plating. Automated imaging and quantification showed that, while the number of SMN+ gems per ISL1+ MN did not change upon the presence of the Clover tag at the earliest time-point (Figure 1G-I), significantly more were detected in the BJ SMN:Clover MNs after 8 days in culture. This may be explained by the higher relative fluorescence of the SMN:Clover+ gems compared to that of the antibody-labeled ones, at both time-points measured (Figure S2A-B), supporting that the SMN:Clover signal is brighter than the antibody-labeled SMN protein and therefore more easily identified by the image analysis script. In order to investigate whether the Clover tag affected the assembly/disassembly of nuclear gems, MNs derived from two BJ SMN:Clover lines (#16 and #23) were treated with Cullin-E3-ligase inhibitor MLN4924 (that prevents SMN proteasomal-mediated degradation (Rodriguez-Muela et al., 2017)) or cycloheximide (CHX, that halts protein synthesis) for 3 days, fixed and immunostained against SMN. The number of endogenously labeled SMN:Clover+ and immunostained SMN+ gems per MN was quantified. As expected, MLN4924 treatment increased the number of Clover+ gems and CHX had the opposite effect (Figure S2C-D). This indicates that the dynamics of the nuclear gems are not impacted by the Clover reporter. Co-immunoprecipitation analysis using anti-SMN or anti-GFP antibodies from BJ parental and the edited BJ clone #23 hiPSC lysates showed that SMN:Clover binds to untagged SMN protein and SMN main binding partner, gemin2, similarly to untagged SMN protein (Figure 1J). This indicates that Clover does not interfere with SMN self-oligomerization and binding with major SMN complex components.

Finally, we measured SMN:Clover protein half-life by treating the parental SMA hiPSC lines and their isogenic corrected clones with CHX, a method previously used to study SMN turnover (Lorson and Androphy, 2000; Vitte et al., 2007; Locatelli et al., 2015) that yielded ∼48 hours in hiPSC-derived MNs (Rodriguez-Muela et al., 2017). Unbiased, automated imaging and quantification of SMN:Clover fluorescence intensity in both corrected clones for the three SMA hiPSC lines showed a half-life of approximately 40 hours (Figure 1K-N). These results indicate that the presence of the fluorescent tag does not significantly alter SMN protein turnover. The same treatment paradigm followed by quantification of immunostained SMN showed a comparable degradation profile for total SMN (untagged and tagged with Clover) of ∼40h half-life in the parental BJ and 39C-III lines and the isogenic clones (Figure 1P and S2E-F). However, the 50% decline in SMN protein levels for the 51N-II and 38D-I parental lines were reached later (Figure 1 O, Q-R and S2G-H), which suggests that cells activate specific pathways for SMN protein regulation when its levels are below a certain threshold. Western blot (Figure 1D-E) and immunostaining-based analysis (Figure 1P-R) also revealed that the corrected hiPSCs have a 3-4 fold increase in SMN protein compared to their parental lines. A comparable decline in the number of cells upon CHX treatment over time was observed between the lines, indicating similar accumulated toxicity (Figure S2I-L). Analogous degradation profile results were obtained for gemin2 protein levels (Figure 2A). Using this method gemin2 showed a ∼45-50h half-life in the BJ WT hiPSCs and the corrected clones, which displayed a degradation pattern parallel to their SMA parental lines. Unlike for SMN protein, the corrected clones did not show increased gemin2 protein levels compared to their parental lines (Figure 2F-H), which indicates that tagging endogenous SMN protein does not alter the levels of this main binding partner. Together, these results indicate that endogenously tagged SMN:Clover displays subcellular localization, binding to key partners, turnover rate and response to modulators of its stability analogous to the untagged SMN fraction.

**Figure 2.**
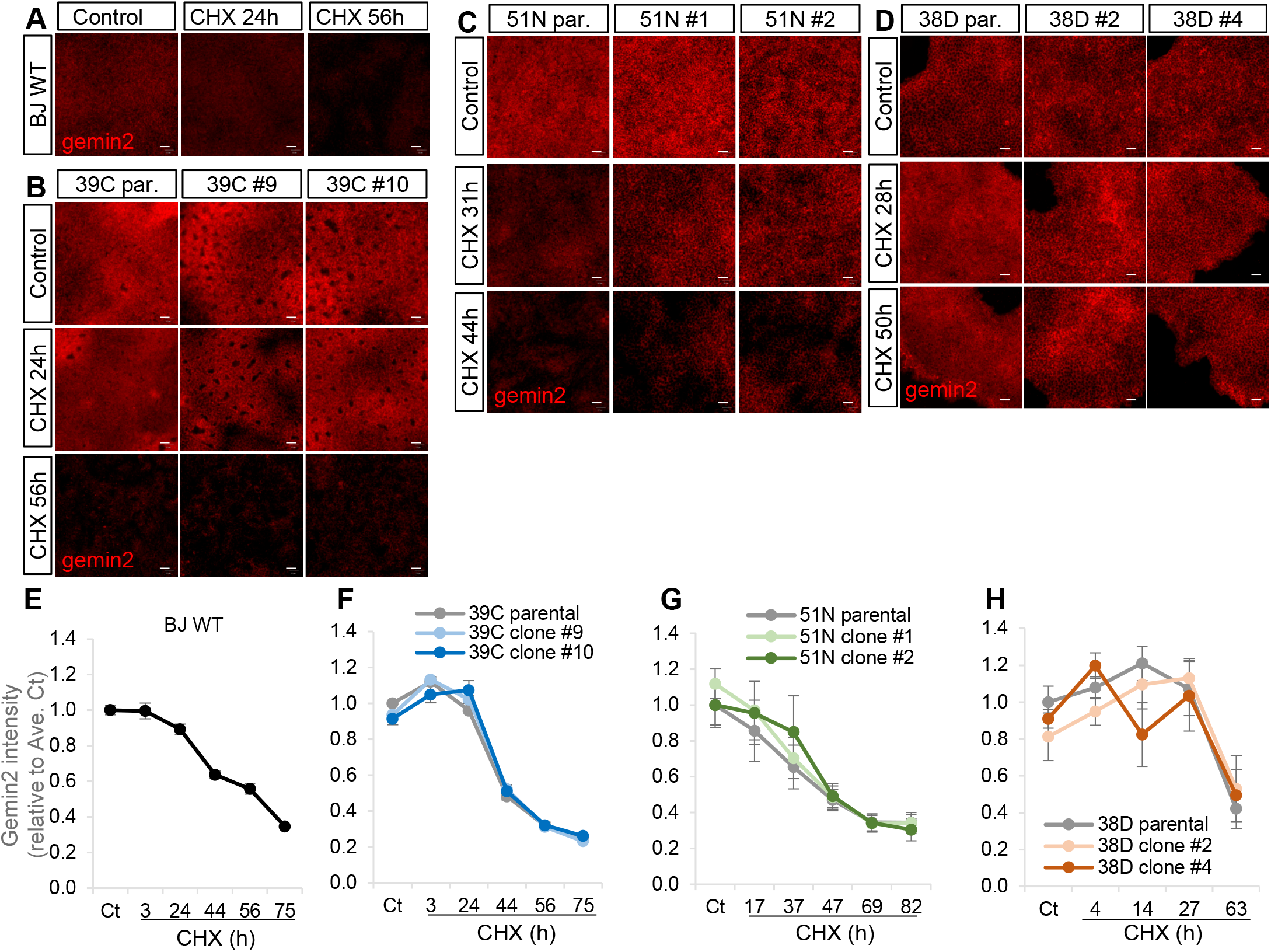
The abundance and degradation rate of the main SMN-Binding partner Gemin2 is not altered by Clover SMN. (**A**) Representative images showing immunostaining against Gemin2 in healthy BJ hiPSCs, (**B**) SMA 39C-III and its corrected clones, (**C**) SMA 51N-II and its corrected clones and (**D**) SMA 38D-I and its corrected clones after treatment with CHX for the indicated times or vehicle (Control, DMSO). Scale bar, 50 µm. (**E-H**) Quantifications of Gemin2 immunofluorescence intensities of the respective lines upon the treatments.

### The corrected isogenic hiPSC lines present similar proliferation rates and differentiation potential to healthy control and SMA parental lines

Several studies have reported alterations in cell cycle genes in SMA models (Ruggiu et al., 2012; Maeda et al., 2014). Other publications have shown conflicting results on reduced (Shafey et al., 2005; Wishart et al., 2010; Li et al., 2013; Grice and Liu, 2011) or increased (Shafey et al., 2008; Luchetti et al., 2015) cell proliferation upon SMN loss depending on the cell type and the percentage of SMN reduction. We therefore investigated whether the isogenic corrected SMA lines, harboring significantly higher SMN protein levels than the parental ones, displayed different growth profiles that could potentially influence their differentiation capabilities.

hiPSC colonies from the BJ control and the three SMA isogenic trios were dissociated and single-cell suspensions plated on Matrigel-coated 96-well plates in mTeSR medium. 2 days after plating SiR-DNA, a cell permeable fluorogenic probe for live cell DNA labeling, was added and the cells imaged. The same fields and wells were imaged 2, 4 and 6 days later and the area occupied by the labeled hiPSC nuclei automatically quantified. No significant difference was observed in the iPSC colony growth rate between the disease and the BJ control lines (Figure 3A, E and S3A-B), although the type I line exhibited an increased vulnerability compared to the other lines, shown by the reduced number of cells that initially attached to the plate (Figure 3E). No significant difference was either observed between the corrected clones and their isogenic SMA counterparts (Figure 3B-E and S3A). The quantification of P-H3, a commonly used mitotic marker, on fixed and immunostained plates at multiple time points after plating also revealed no robust difference between the SMA parental and the corrected clones (data not shown). Therefore, no morphological or growth abnormalities were detected in the SMA parental hiPSCs compared to BJ WT or between the isogenic corrected hiPSCs and the parental SMA ones. These data align with the argument that SMN’s control of cell cycle progression and cell proliferation is highly cell-type specific.

**Figure 3.**
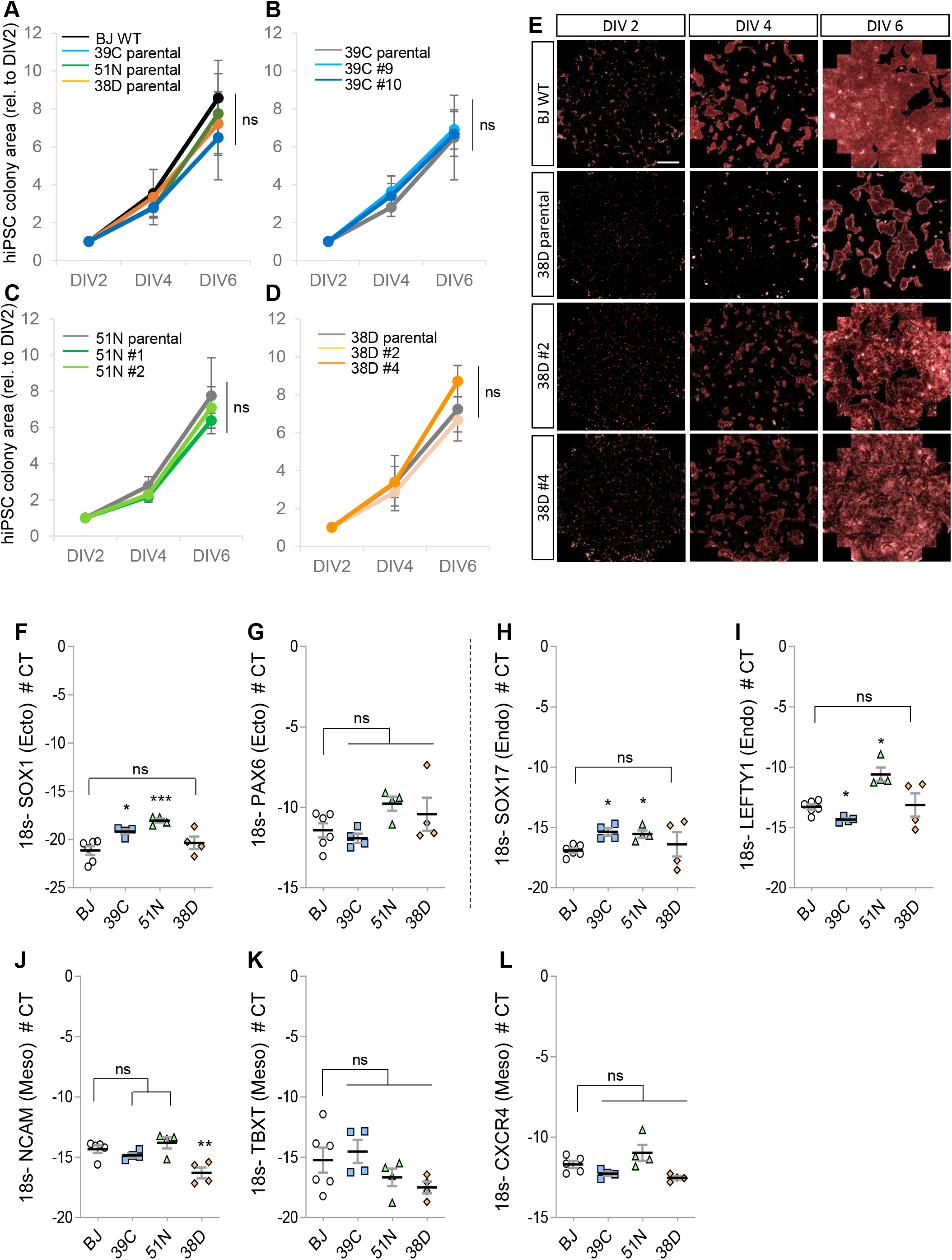
The corrected isogenic SMA hiPSCs proliferate and differentiate into the three-germ layer similarly to the SMA parental lines and healthy control. Quantification of the hiPSCs colonies growth over time. The graphs represent the sum of the area (µm^2^) occupied by the hiPSC colonies in the 96w plates, overtime and relative to the first time point measured for each of the lines. The whole area of 10 wells in each 96w plate was imaged and quantified. Colony area growth of BJ WT compared to the 3 SMA lines (**A**), 39C-III compared to its 2 isogenic clones (**B**), 51N-II compared to its 2 isogenic clones (**C**) and 38D-I compared to its 2 isogenic clones (**D**) (Two-way ANOVA followed by Tukey’s multiple comparison test, n=3-4). (**E**) Representative images showing SiR-DNA-labeled hiPSC nuclei from the BJ WT and SMA 38D-I and its corrected clones after being plated for 2, 4 and 6 days. Scale bar, 1 mm. qPCR quantification of ectodermal *SOX1* (**F**) and *PAX6* (**G**), endodermal *SOX17* (**H**) and *LEFTY1* (**I**) and mesodermal *NCAM* (**J**), *TBXT* (**K**) and *CXCR4* (**L**) markers from BJ WT and SMA hiPSCs cultured in STEMdiff™ Trilineage Differentiation Kit (see Materials and Methods). Graphics represent CT mRNA expression levels of the indicated genes subtracted to the housekeeping gene (18s) expression levels (*p<0.05, **p<0.01, ***p<0.005 One-way ANOVA followed by Tukey’s multiple comparison test, n=4).

Different hiPSC lines have various predisposition to differentiate into the 3-germ lineages. To rule out that the genome editing introduced an undetected off-target effect which could potentially alter the differentiation capabilities of the lines, we performed the next assays. The differentiation capacity of both the BJ WT and the three SMA lines has been separately published before (Ng et al., 2015; Rodriguez-Muela et al., 2017). To do a side-by-side comparison, the same number of hiPSCs from the 4 lines were plated and cultured in STEMdiff™ Trilineage Differentiation media. To assess the hiPSC differentiation potential, we measured the expression levels of *SOX1* and *PAX6* (ectodermal markers), *TBXT, NCAM* and *CXCR4* (mesodermal markers) and *SOX17* and *LEFTY1* (endodermal markers) genes by qPCR. First, using the BJ WT line, we validated that the expression of those genes was highest in the hiPSCs cultured with the corresponding differentiation medium (Figure S3C-I). Next, we measured the expression levels of those 7 genes in the BJ WT and the 3 SMA parental hiPSCs cultured with the three different media. While significant differences were occasionally detected between the control line and one or two SMA lines (e.g. 39C-III and 51N-II showed higher *SOX1* levels than BJ), no consistent bias for any of the SMA lines towards a specific lineage was detected under these culture conditions (Figure 3F-L). Finally, we asked whether our corrected isogenic SMA lines showed any major difference with their parental disease lines in their differentiation potential. Similarly, we did not detect robust differences in the expression levels of the lineage-specific genes measured between the corrected clones and their isogenic parental SMA lines (Figure S3J-L). Together, these results demonstrate that the isogenic corrected hiPSCs do not exhibit major differences in proliferation or 3-germ layer differentiation potential compared to the parental lines.

### The progressive death of SMA MNs is ameliorated in the corrected isogenic lines

Although multisystemic abnormalities have been widely reported for SMA in the last decade (Lipnick et al., 2019; Yeo and Darras, 2020), MNs are the cell type most affected by an SMN deficiency, and the molecular underpinnings of such selective vulnerability remain unclear. Others and we have shown that SMA hiPSC-derived spinal MNs die faster in culture than the ones not carrying *SMN1* mutations (Ng et al., 2015), and that increasing SMN protein levels promotes MN survival (Rodriguez-Muela et al., 2017). However, to our knowledge, these assays have not been performed in an isogenic context. To investigate whether in our isogenic SMA model MN death is mitigated, we subjected the healthy control line and the three isogenic trios to an EB-based MN differentiation protocol that we have previously used (Rodriguez-Muela et al., 2017; Rodriguez-Muela et al., 2018) (Figure S4A). At the end of the differentiation protocol the spinal cord spheres were dissociated and the neurons plated. To avoid the variability intrinsically bound to quantifying cell numbers across different plates, the same MN culture plate was stained with SiR-DNA 2 days after plating and imaged the same day and 10 days later. The quantification of the number of living neurons after 12 days for each line relative to how many were present at day 2 showed that only 60% of the 51N-II and 38D-I SMA MNs survived in that period compared to the survival of the healthy control line (Figure 4A). SMA 39C-III MNs did not show a significant reduction in their survival compared to BJ WT (Figure 4A). Importantly, both isogenic clones for each of the significantly less fit lines showed a significantly higher MN survival rate than the disease parental. The degree of MN rescue corresponded to the severity of the line. While the 39C isogenic clones (#9 and #10) only showed a not-significant 20-30% increased survival over the parental SMA 39C-III MNs (Figure 4C and S4B), ∼40% was the increased survival for the isogenic 51N-II MNs (Figure 4C and S4C) and 50-70% for the isogenic 38D-I lines (Figure 4D-E). These results support that the more severe the pathological phenotype is (e.g. more neuronal death overtime), the greater the potential rescue.

**Figure 4.**
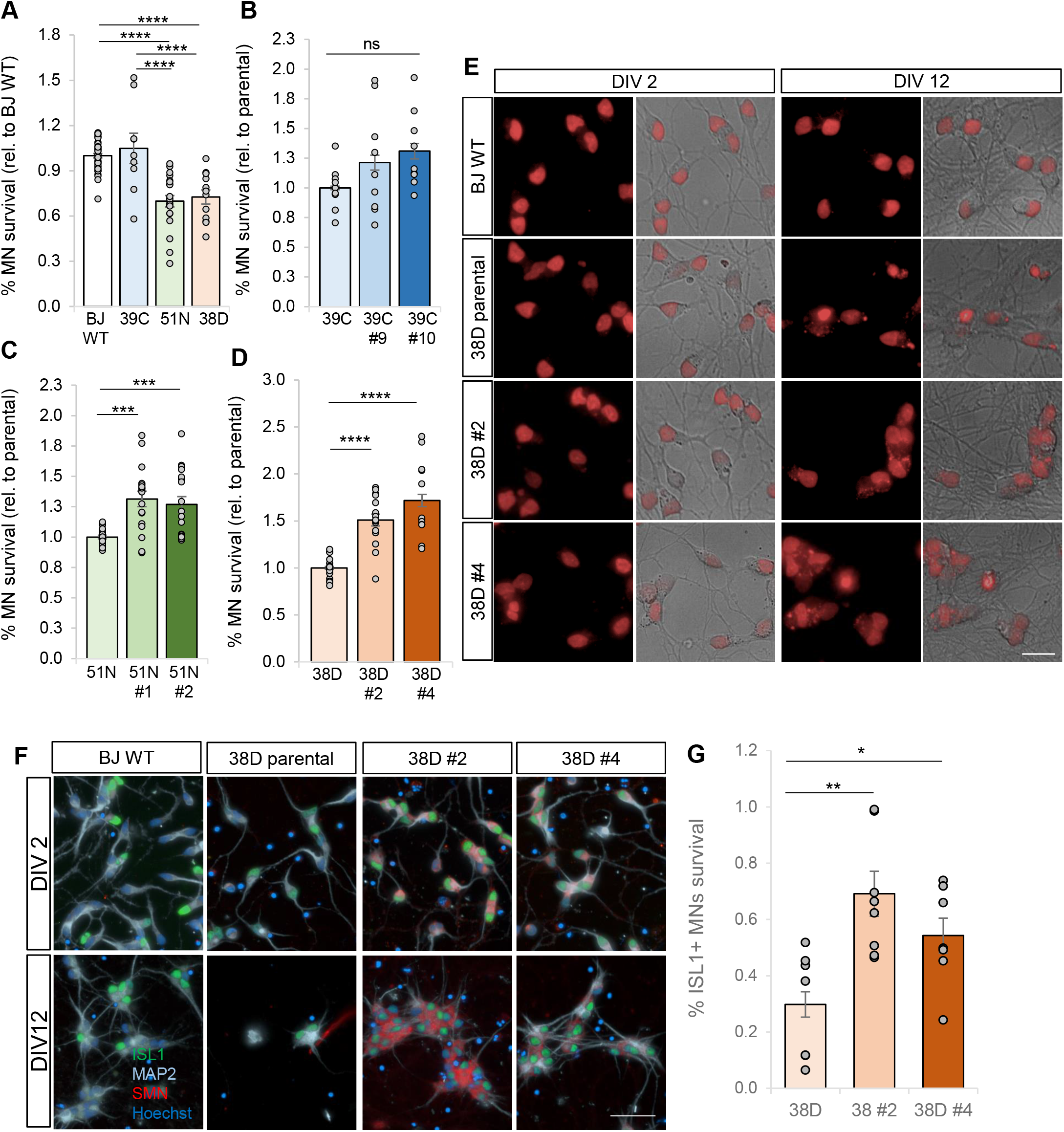
SMA hiPSC-derived spinal MN death is ameliorated in the isogenic corrected lines. Quantification of the percentage of hiPSC-derived neurons that survive 10 days after being plated in 96w plates. Two days after plating the MN cultures were labeled with SiR-DNA and imaged live (Operetta CLS). The same fields and wells were imaged again after 10 days and the percentage of surviving neurons quantified. The graphs show these values for the 3 SMA lines relative to the BJ WT (**A**), the percentage survival of the type III isogenic corrected cultures relative to the 39C-III parental (**B**), the type II isogenic corrected cultures relative to the 51N-II parental (**C**) and the type I isogenic corrected cultures relative to the 38D-I parental (**D**). Each dot in the graphs represents individually analyzed wells of all conducted experiments (***p<0.001, ****p<0.0001 One-way ANOVA test followed by Tukey’s analysis, n=4-7 independent experiments per line). (**E**) Representative images of BJ WT, SMA type I and both isogenic corrected iPSC-derived MN cultures stained with SiR-DNA (red) after 2 and 12 days in culture. Bright field images of the corresponding neurons are also shown. Scale bar 20 µm. (**F**) Representative images of BJ WT, SMA 38D-I and both isogenic corrected hiPSC-derived MN cultures fixed and immunostained after 2 and 12 days in culture against ISL1 (green), MAP2 (cyan) and SMN (red) antibodies. Nuclei are stained with Hoechst (blue). Scale bar 50 µm. (**G**) Quantification of the percentage of ISL1+ MNs derived from the SMA type I and the isogenic corrected clones that survived after 12 days compared to the number quantified after 2 days in culture (p <0.05, **p<0.01One-way ANOVA test followed by Fisher’s LSD analysis, n=7).

Next, to confirm that the SiR-DNA live-imaged cells were indeed neurons and that a large percentage of them expressed MN markers, dissociated spinal cord spheres generated from the type I isogenic trio were fixed and immunostained after 2 and 12 days in culture. Over 95% of the cells were MAP2+ neurons and ∼60% expressed ISL1 (Figure S4D-E). Using this approach to determine MN survival, only ∼30% of the SMA 38D-I MNs survive 12 days in culture, whereas the isogenic clones showed a ∼60% survival rate (Figure 4F-G). Together, these results indicate that correcting the *SMN2* mutation, and therefore significantly increasing the amount of SMN protein, notably ameliorated MN death, the main SMA pathological hallmark. Interestingly, despite the isogenic corrected MNs have a 3-4 fold increase in SMN protein levels over their disease counterparts, this phenotypic rescue is nevertheless not complete. This unexpected result could be due to SMN-independent stress-induced cell death occurring during the sphere dissociation process or intrinsic to the culture conditions. It is also possible that factors other than SMN deficiency contribute to the MN death in SMA (see discussion). In such case the use of isogenic models to tackle disease-specific mechanisms, rather than utilizing hiPSC lines from healthy donors to compare to the disease ones, constitutes a better approach to generate disease-relevant knowledge.

### SMA hiPSCs show an impaired spinal cord organoid formation and the corrected isogenic lines partially restore that phenotype

Once demonstrated that our isogenic model recapitulates the most important SMA hallmark, MN death in a disease severity dependent manner, and that the mutation-corrected clones showed a significant attenuation of such phenotype, we aimed to shed light on the unresolved question of how SMN protein is essential for the survival of MNs specifically. To do so, we developed a spinal cord organoid (SCO) protocol to study MN specification, maturation and survival using our cohort of isogenic SMA lines.

We adapted our EB-based differentiation protocol to generate a more robust model in which we also included an extracellular matrix. Briefly, 4.000 cells from the 10 lines (BJ WT and the 3 isogenic trios) were seeded per well into ULA 96w plates and cultured in mTeSR for 5 days. The hiPSCs self-assembled into a main cell aggregate or sphere that was detectable 16 hours later in an extremely robust manner (Figure S5A). The spheres were changed to neural induction medium, embedded in matrigel drops and exposed to small molecules to induce their differentiation towards ventral spinal cord (described in detail in the Materials and Methods section) (Figure 5A). First, we imaged the ULA plates and determined the percentage of stem-cell spheres that self-assembled 2 days after being transferred into the ULA 96w plate and the percentage that assembled after a week in culture, before the spheres were too large to be fully captured by a 10X objective. While 100% of the wells where BJ WT hiPSCs were seeded showed a small sphere (∼150 µm in diameter) (Figure 5B, F), only 40% of the most severe SMA line 38D-I assembled into much smaller and irregular shapes (Figure 5E, F). By completion of the imaging assay this value reached 70% in the 38D-I. The least severe line, 39C-III, showed a ∼60% average sphere formation rate and by completion the stem cells in all wells had self-assembled (Figure 5C, F). Surprisingly, the SMA line 51N-II, despite carrying the same *SMN2* copy number (3 copies (Rodriguez-Muela et al., 2017)) as the type III, and not having significantly different SMN protein levels (Rodriguez-Muela et al., 2017), did not exhibit a defect, although the spheres were initially smaller than the BJ WT ones (Figure 5D, F). The two corrected type I clones partially fixed the self-assembly deficiency displayed by the parental disease line, showing by initiation 60-80% sphere formation rate and by completion 80-100%, for clones #4 and #2 respectively (Figure 5E, J). Both 39C corrected clones showed full sphere formation efficiency already at the initiation of the assay (Figure 5C).

**Figure 5.**
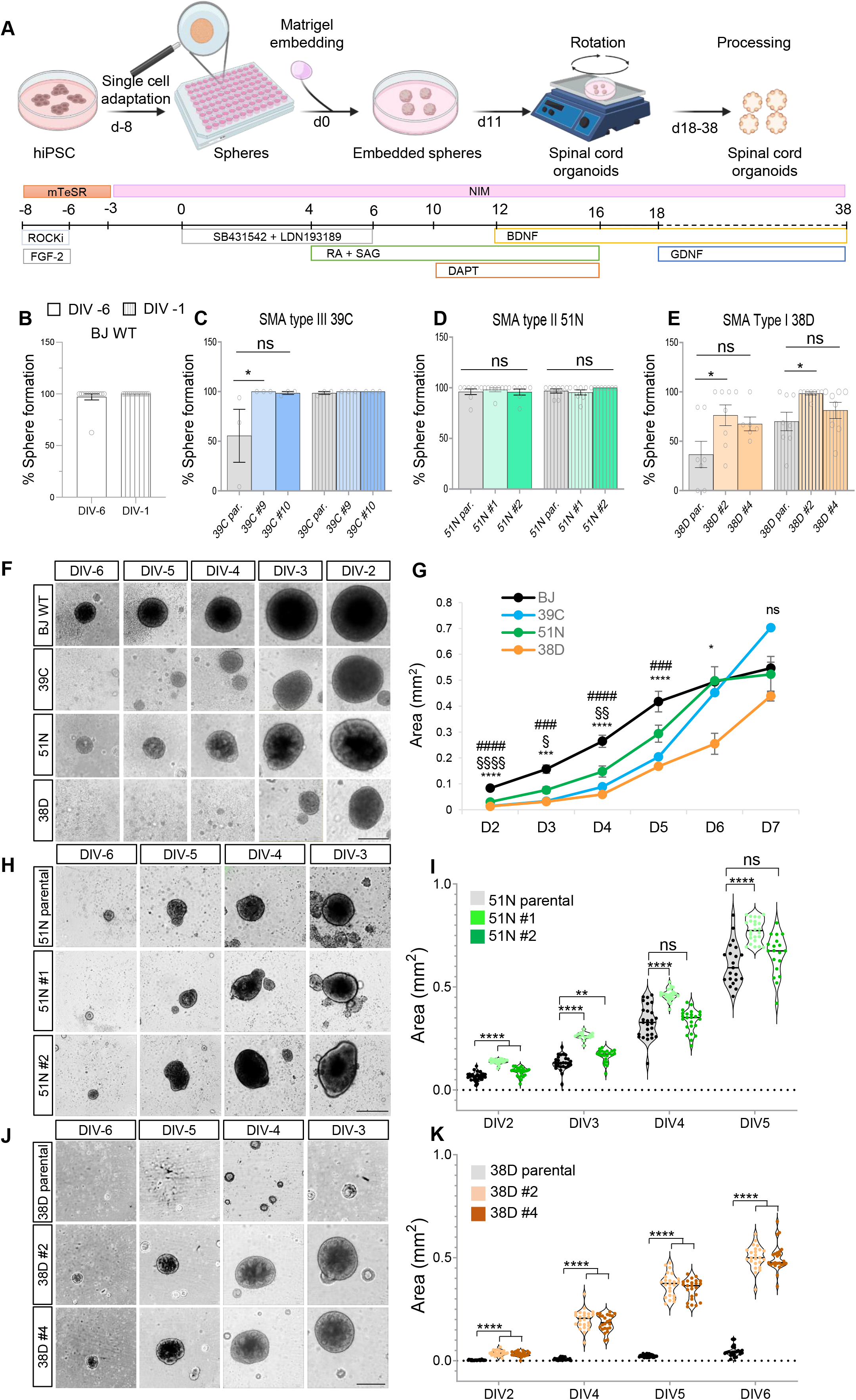
SMA hiPSCs present a defective sphere formation and growth when subjected to a spinal cord organoid formation protocol, which is improved the isogenic corrected lines. (**A**) Schematic representation of the protocol followed to generate spinal cord organoids from the isogenic SMA and corrected hiPSC lines. Quantification of the percentage of hiPSC-seeded wells that showed a sphere two days after the hiPSCs were seeded as a single-cell solution and after 7 days in culture for the healthy control BJ WT line (**B**), SMA 39C-III line and its isogenic corrected clones (**C**), 51N-II and corrected clones (**D**) and 38D-I and corrected clones (**E**) (Kruskal-Wallis analysis followed by Dunn’s test, *p<0.05, n=3-6). (**F**) Representative images of the self-assembled spheres generated from the BJ WT and SMA parental hiPSCs at the indicated times after being seeded as a single-cell solution in ultra-low attachment 96w plates. Scale bar 400 µm. (**G**) Quantification of the BJ WT and SMA hiPSC-derived sphere area (mm^2^) increase over time. The graph represents the average of at least 30 spheres imaged and quantified per line and per experiment over the indicated time course (*p<0.05, ***p<0.001, ****p<0.0001 Two-way ANOVA test followed by Tukey’s analysis; # represents comparisons between BT WT and 39C-III; § between BT WT and 51N-II and * between BT WT and 38D-I, n=4-8). (**H**) Representative images of the self-assembled spheres generated from the SMA parental 51N-II line and the isogenic corrected clones #1 and #2 at the indicated times after being seeded as a single-cell solution in ultra-low attachment 96w plates. Scale bar 400 µm. (**I**) Quantification of the area (mm^2^) of SMA 51N-II and isogenic corrected hiPSC-derived spheres 2 days after the hiPSC were seeded and until day 5. The violin plot exemplifies the size distribution of the individual spheres quantified for one representative experiment (**p<0.01, ****p<0.0001 Two-way ANOVA test followed by Tukey’s analysis, n=30 spheres). (**J**) Representative images of the self-assembled spheres generated from the SMA parental 38D-I line and the isogenic corrected clones #2 and #4 at the indicated times after being seeded as a single-cell solution in ultra-low attachment 96w plates. Scale bar 400 µm. (**K**) Quantification of the area (mm^2^) of SMA 38D-I and isogenic corrected iPSC-derived spheres 2 days after the hiPSCs were seeded and until day 6. The violin plot exemplifies the size distribution of the individual spheres quantified for one representative experiment (****p<0.0001 Two-way ANOVA test followed by Tukey’s analysis, n=30 spheres).

The quantification of the average sphere area over this period confirmed that BJ WT spheres were the largest from the beginning of the differentiation protocol, the 38D-I the smallest and the 51N-II displayed the mildest phenotype in this assay (Figure 5F-G). The isogenic types III and I clone-derived spheres were significantly larger than their corresponding parental ones (Figure 5J-K and Figure S5B). Interestingly, although two of the three SMA hiPSCs showed this robust sphere self-assembly defect, the spheres that did assemble and survived throughout the culture grew at a similar rate or even faster than the BJ WT (Figure S5C-F). This suggests that these SMA hiPSCs do not present a general growth failure, but rather an impaired capability of self-assembling and/or surviving into a stem cell aggregate. To validate this aspect, an increasing number of hiPSCs, from 600 to 20.000 cells, from the BJ WT and the SMA lines were seeded into ULA 96w plates and imaged 5 and 15 days later. While 600 hiPSCs was enough for the BJ WT line to robustly self-assemble into one uniform small sphere, the 38D-I line required up to 5.000 cells (Figure S5F-G). 600 cells were also enough for the 39C-III line to self-assemble although not robustly. In addition, increasing the number of seeded cells for this line did not lead to an improved sphere formation but to multiple small and aberrantly shaped spheres, most of which died, and the remaining ones fused into a large and amorphous cell aggregate (Figure S5F-G). The 51N-II line similarly to the BJ WT, produced a single, healthy-looking sphere in the 600 cell-seeded wells, however, these were smaller than the BJ ones (Figure S5F-G). Altogether, these results indicate that SMA hiPSCs present a deficient capacity to self-assemble into stem cell aggregate structures when plated as a low concentration solution of dissociated cells and the isogenic corrected clones notably ameliorate that phenotype.

### SMA SCOs show altered neurodevelopment, which is only partially corrected in the isogenic control lines

While there has been some controversy on whether or not SMA motor axons showed a defect in formation and outgrowth during development (Mcgovern et al., 2008; Hao Le et al., 2013; Hao Le et al., 2017; Motyl et al., 2020; Kong et al., 2021), several recent studies have described abnormalities in MN progenitors resulting in defective MN migration, target innervation and survival (Park et al., 2010; Grice and Liu, 2022; Simic et al., 2008). Neurogenesis defects in several brain areas have also been reported in SMA model organisms (Wishart et al., 2010). These findings together with the observed impairment in self-organization of our SMA hiPSCs into stem cell aggregates and reduced MN survival, led us to investigate how SMN deficiency could impact the formation of neural progenitors, neurogenesis and MN differentiation in our isogenic human system.

SCOs from the healthy BJ WT line and the most severe SMA 38D-I line, together with its isogenic corrected clones were generated following the protocol described above (Figure 5A) and 8 days after being embedded in matrigel they were processed for mRNA expression or fixed for immunostaining analysis of neural stem cell (NSC) markers. Interestingly, the mRNA expression level of *SOX2* and *NESTIN* was significantly reduced in the SMA 38D-I line compared to the BJ WT (Figure 6A-B). *NGN2* mRNA levels, a pro-neural gene positively regulated by *SOX2 (Amador-Arjona et al., 2015)*, was, accordingly, significantly downregulated in the disease line (Figure 6C). The isogenic corrected clones partially corrected that phenotype for the three genes studied (Figure 6A-C). Immunostaining of SCO cryosections confirmed these results (Figure 6D-E, S5H) and, notably, uncovered the presence of NESTIN+ cells that were negative for SOX2 in the SMA organoids (Figure 6D’). The expression of both NSC genes, *SOX2* and *NESTIN*, during CNS development is commonly found concomitant; however, subpopulations of progenitors expressing only one of the markers have been described in a neurodegeneration context before (Albright et al., 2016). It is therefore possible that specific NSC subpopulations are altered in the spinal cord upon SMN deficiency, which might result in abnormalities in the lineage specification of certain neuronal types later in development.

**Figure 6.**
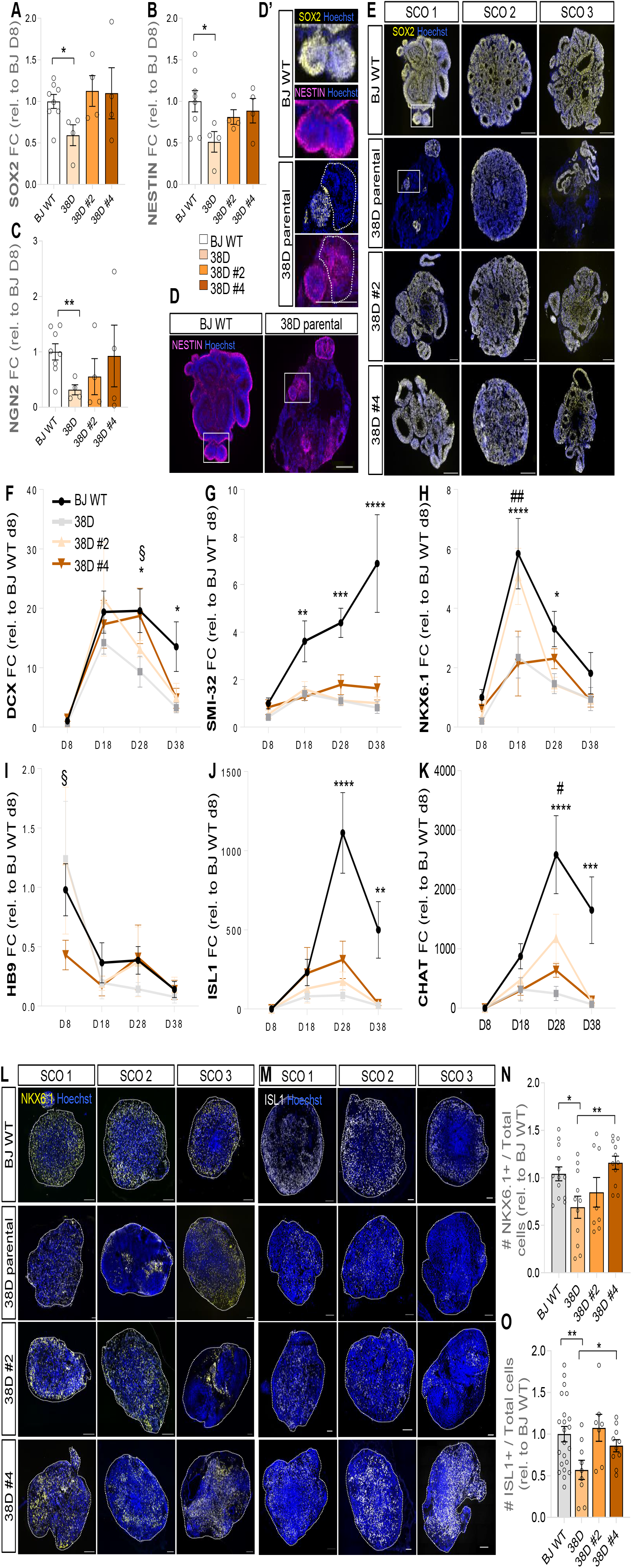
SMA spinal cord organoids show signs of altered neural development, a phenotype that is partially corrected in the isogenic controls. qPCR showing SOX2 (**A**), NESTIN (**B**) and NGN2 (**C**) mRNA expression in spinal cord organoids (SCO) derived from BJ WT, SMA 38D-I and both isogenic corrected clones 8 days into the differentiation protocol. Gene expression is indicated as fold change of 2-ΔΔCt with respect to 18s, normalized to BJ WT SCO (*p<0.05, **p<0.01 One-way ANOVA followed by Fisher’s LSD multiple comparison test, n=4, 4-8 pooled SCOs per experiment). (**D**) Representative images showing BJ WT and severe SMA 38D-I day 8 SCOs after being fixed, cryosectioned and immunostained against NESTIN (magenta; nuclei stained with Hoechst, blue). Scale bar 100 µm. The white squares represent magnified areas displayed in (**D’**) where SOX2 (yellow) is also shown. The region surrounded by dotted line indicates a SCO area with presence of NESTIN signal and depleted from SOX2. Scale bar 100 µm. (**E**) Representative images from BJ WT, 38D-I and isogenic corrected day 8 SCOs showing SOX2 (yellow). Nuclei stained with Hoechst, blue. Scale bar 100 µm. Note that BJ WT and SMA 38D-I parental “SCOrg 1” show in D-D’ and E are the same. qPCR showing doublecortin (DCX) (**F**), SMI-32 (**G**), NKX6.1 (**H**), HB9 (**I**), ISL1 (**J**) and CHAT (**K**) mRNA expression in SCO derived from BJ WT, SMA type I 38D and both isogenic corrected clones 8, 18, 28 and 38 days into the differentiation protocol. Gene expression is indicated as fold change of 2-ΔΔCt with respect to 18s, normalized to day 8 BJ WT SCO (*p<0.05, **p<0.01, ***p<0.005, ****p<0.0001 Two-way ANOVA followed by Fisher’s LSD multiple comparison test, n=4, 4-8 pooled SCOs per experiment. * represents difference between BJ WT and 38D-I; # between 38D and 38D clone #2 and § between 38D and clone #4). (**L**) Representative images showing BJ WT, SMA 38D-I and corrected day 28 SCOs fixed and immunostained against NKX6.1 (yellow) and ISL1 (white) (**M**). Scale bar 100 µm. (**N**) Quantification of the number of NKX6.1+ cells relative to the total number of cells (stained with Hoechst) in day 28 SCO from BJ WT, SMA 38D-I and isogenic corrected clones. (**O**) Quantification of the number of ISL1+ cells relative to the total number of cells (stained with Hoechst) in day 28 SCO from BJ WT, SMA 38D-I and isogenic corrected clones (*p<0.05, **p<0.01 One-way ANOVA followed by Fisher’s LSD multiple comparison test, n=3-6, 3-4 SCOs per experiment).

Considering these findings, we were next interested in exploring whether the expression of pan-neuronal and MN specific genes was altered in the disease SCOs and corrected in the isogenic clones. Time-course gene expression analysis at day 8, 18, 28 and 38 of the differentiation protocol was performed. Importantly, using BJ WT SCOs as reference, we observed that the expression levels of all genes studied changed throughout this period as expected according to the well-known genetic programs that govern spinal cord developmental *in vivo* (Sagner and Briscoe, 2019). Specifically, the levels of the neuronal migration protein doublecortin (*DCX*), often used as immature neuronal marker, drastically increased from day 8 to day 18 and declined after day 28, as expected when neurons start to mature and lose their expression (Figure 6F). The pan-neuronal marker *SMI-32* gradually increased during the differentiation process (Figure 6G). The expression of the spinal cord MN and ventral V3 progenitor domains *NKX6.1* peaked at day 18 to later decrease (Figure 6H). The expression of the homeobox gene *HB9*, a terminal MN transcription factor in the developing spinal cord, was highest at the first time-point measured, day 8, and decreased from then onward (Figure 6I). *HB9* expression declines during development (Heindorf et al., 2019; Arber et al., 1999) and longitudinal gene expression studies revealed that it begins to appear 3-5 days after addition of patterning molecules in mouse EB-based ESC-derived MN differentiation protocols (Wichterle et al., 2002; Soundararajan et al., 2006; Amoroso et al., 2013) and 7-14 days in human EB-based iPSC-derived protocols (Amoroso et al., 2013). Our *HB9* expression results indicate that in our SCO protocol the specification of MN identity occurred at an early stage. The expression of another canonical MN identity transcription factor, *ISL1*, and a more mature cholinergic MN marker, *CHAT*, gradually and markedly increased from day 8, peaking at day 28 (Figure 6J-K). The observed decline in the expression of these two markers from day 28 to 38, together with the increasing SMI-32 levels, could be explained by either the death of MNs at this latest stage or by the emergence of other neuronal types, lowering the relative expression of MN markers.

The comparison of the SMA type I SCOs to the healthy SCOs revealed that *DCX* expression was significantly reduced at all time-points (Figure 6F). Similar results were observed for *SMI-32* (Figure 6E). Interestingly, while *DCX* expression profile was increased in the isogenic corrected clones compared to the disease parental, *SMI-32* expression remained as low in the clones as in the SMA SCOs. The expression of *NKX6.1* was also significantly lower in the SMA SCOs than in the BJ WT at all time-points studied (Figure 6H), as well as *HB9* from day 18 to 38 (Figure 6I), *ISL1* (Figure 6J) and *CHAT* (Figure 6K). Interestingly, the last two genes reached highest expression levels at day 18 in the SMA SCOs instead of at day 28 as in the WT line, pointing to an accelerated MN differentiation in the disease SCOs. This phenomenon was reverted in the isogenic corrected SCOs. The expression of these four MN progenitor and postmitotic MN markers in the isogenic corrected clones showed an intermediate profile between the healthy control and the SMA SCOs for most time-points. Notably, however, *ISL1* and *CHAT* expression dropped in the corrected SCOs closer to disease values by day 38. As mentioned above, a similar decline was observed in the BJ WT SCOs relative to their day 28 higher expression values. However, unlike for the BJ WT, the constant SMI-32 levels expressed in the SMA and the corrected SCOs suggests that MNs do die in the SMA SCOs and that the rescue effect detected in the corrected SCOs is not maintained beyond a certain point.

Immunostaining analysis on cryosections confirmed a reduction in the number NKX6.1+ progenitor cells in day 28 SMA SCOs compared to the healthy controls and their numbers markedly increased in the isogenic SCOs (Figure 6L, N). The number of ISL1+ cells in day 28 38D-I SCOs was also significantly less abundant compared to the healthy BJ WT and a marked increase in the corrected SCOs was detected (Figure 6M, O). Similar findings were observed for the pan-neural marker MAP2 (Figure S5I). Interestingly, the number of ISL1+ MNs in day18 SMA SCOs was higher than at day 28, whereas the opposite was noticed for the BJ WT and the isogenic corrected SCOs (Figure S5J-K), in agreement with our gene expression analysis and reinforcing the hypothesis of an abnormal MN specification timeline in SMA.

Together these results show that the isogenic control SCOs displayed a developmental pattern that generally resembled the one expected and observed from non-disease SCOs in terms of time and relative expression of NSC, progenitors and MN markers. However, despite having notably higher SMN amounts than their SMA counterparts, several disease phenotypes were not corrected (e.g. defective progressive neuronal emergence and maturation, depicted by SMI-32, or drastic loss of canonical MN markers at a later developmental stage). Furthermore, it is crucial to remark that, although a healthy control line -used in numerous studies before (Ng et al., 2015; Rodriguez-Muela et al., 2017; Rodriguez-Muela et al., 2018; Paik et al., 2018; Wu et al., 2019; Rigamonti et al., 2016; Hor et al., 2018; Maherali et al., 2008)-was employed to define a normal developmental baseline, it evidently has a different genetic background to the SMA lines. Therefore, the most relevant biological comparisons are the ones made between each of the SMA lines and their isogenic corrected controls.

## Discussion

How the disturbance of a basic homeostatic function (mRNA splicing) leads primarily to a neuromuscular disorder remains puzzling. It is now clear that different cell types present a distinct sensitivity to the same reduction in SMN protein levels. It is also generally accepted that spinal MNs are especially vulnerable to a SMN deficiency, and much effort has been devoted to identifying the MN-specific functions of SMN that could explain that selective vulnerability. Yet, there is still no consensus on whether this selectivity is solely due to cell intrinsic abnormalities or whether cell non-autonomous mechanisms play a central role, let alone whether developmental marks render those neurons to degenerate months or years after having developed. In this study, we hypothesized that the MN selective vulnerability that characterizes SMA is already imprinted during their early development. We generated the first cohort of isogenic hiPSC lines and employed a hiPSC-derived spinal cord organoid (SCO) model to investigate this postulate.

To study human disease using patient-derived hiPSCs models, an isogenic context is essential to uncover disease-relevant mechanisms, otherwise hidden or even misinterpreted when comparing cells obtained from different donors with different genetic backgrounds. More and more studies are now utilizing isogenic pairs as the best control when using hiPSCs to model disease, however, SMA’s complex genetics have hindered the development of this asset. Localized in an unstable region of chromosome 5q (11.1–13.3), that contains a large 500 kb inverted repeat element, *SMN* genes comprise 62% interspersed repetitive DNA and the density of Alu elements are 4-fold higher than in average in the genome (Rochette et al., 2001; Smit, 1999). DNA rearrangements in this region might be the reason why genome editing approaches to correct the exon 7 *SMN2* SMA-causing mutation have proven challenging. Taking advantage of a cloning vector that we have newly generated and following a CRISPR/Cas9-based approach, we have successfully produced several corrected hiPSC clones from three different SMA hiPSC lines. To our knowledge, this is the first isogenic SMA model, generated from multiple SMA hiPSC lines, using a CRISPR/Cas9 method. An additional unique aspect of our isogenic cohort is that the corrected hiPSCs express full-length SMN tagged in the C-terminus with the fluorescent Clover protein. Introducing this reporter not only drastically facilitated the identification of successfully edited cells, but also allowed an accurate detection and quantification of the protein levels and its subcellular localization in living cells. Through a variety of assays, we have established that Clover does not modify the biology of SMN. We believe that this approach holds clear advantages over overexpression experiments, where the integration of unknown gene copy numbers or their localization in the genome is undetermined.

Neurodevelopmental abnormalities are starting to arise as potentially determinant contributors to pathology in neurodegenerative diseases (Barnat et al., 2020). Studies in SMA animal models and human tissue have revealed that reduced levels of SMN results in the impairment of perinatal development of several brain regions (Wishart et al., 2010; Motyl et al., 2020) and developmentally immature MN axons (Kong et al., 2021). Further, although the three approved drugs for SMA (Reilly et al., 2022) have effectively changed the disease course, there is a high variability in the patient’s response to the treatment and none of them constitutes a cure, particularly if an individual has only 2 *SMN2* copies (Strauss et al., 2022). The defective response of some of these patients could be due to an existing great damage by the time the therapy is applied, but also to a MN degeneration priming during development. Understanding, therefore, when and in which cell type the disease first manifests remain a challenge that, once overcome, should enable the optimization of therapies and even their tailoring to specific phases or types of the disease.

In recent years, several studies describing the cellular composition of the developing and adult human spinal cord have been published (Rayon et al., 2021; Zhang et al., 2021), providing excellent resources to investigate the molecular basis of MNDs. Numerous protocols to generate SCOs from hiPSCs are also now available (Lippmann et al., 2015; Mouilleau et al., 2021; Ogura et al., 2018). While they have been mainly used to tackle different aspects of human spinal cord development, they are still rarely employed to model and interrogate SMA pathology. Interestingly, Varderidou-Minasian and colleagues showed that, by comparing their mass spectrometry (MS) data with that of three MS datasets previously published by different groups on human and mouse SMA tissues, the human proteomic profile comparisons shared a larger overlap than those between the two species, even when the comparisons were made between different tissues (Varderidou-Minasian et al., 2021). This reinforces the principle of using human-derived models to study human disease.

The SCO model that we have developed facilitated the discovery of abnormal NSC emergence and deficient and time-shifted expression of neuronal and MN markers in SMA. This is consistent with previous evidence showing altered protein expression profiles in SMA hiPSC-derived MN cultures during neuralization stage (Varderidou-Minasian et al., 2021). In this study, we have observed that the expression of *SOX2* during the SCO NSC expansion phase was significantly reduced in SMA compared to the healthy SCOs. The levels of another NSC marker, *SOX1*, were not detected as altered in the only report on SMA SCO published yet (Hor et al., 2018). However, this discrepancy may well be explained by the different developmental time-points studied, significantly earlier in our case. The expression levels of other NSC marker genes, *NGN2* and *NESTIN*, were also significantly lower in the SMA SCOs than in the healthy ones. NESTIN, normally localized in the periphery of the SCOs with apical-out and basal-in polarity at an early developmental stage, showed a patchy pattern in the SMA SCOs, which was partially corrected in the isogenic clones. Interestingly, in the SMA SCOs, we found NESTIN+/SOX2-cells associated to ventricle-like structures or their proximity. This finding was highly unexpected as both NSC markers are normally expressed together. Nonetheless, there is evidence of *NESTIN+/SOX2-* progenitor cell populations driving dopaminergic neurogenesis in the adult mouse substantia nigra (Albright et al., 2016). This raises the exciting hypothesis of specific neural progenitor pools being affected in SMA. There is another potential explanation for this presumptive inconsistency. NESTIN is cytoskeletal intermediate filament protein (Hensel and Claus, 2018). Numerous studies have reported defects in cytoskeleton-linked signaling pathways in SMA. Hence, it is also possible that SMN deficiency results in altered NESTIN levels by other means different from the ones that caused the SOX2 dysregulation.

Further, by performing a longitudinal gene expression analysis, and using a healthy control line as development standard, we have identified abnormal expression levels of all neuronal markers studied in the SMA SCOs compared to healthy controls. This is in agreement with previous studies showing reduced DCX levels in SMA mouse brains (Wishart et al., 2010) and deficient TUJ1 and UCHL1 protein levels from SMA and control hiPSC-derived MNs (Fuller et al., 2015). In our study, we have additionally observed that the expression of classical MN markers, ISL1 and CHAT is accelerated in the SMA SCOs. This could indicate an aberrant MN differentiation resulting in the quicker degeneration of postmitotic MNs that we have also observed via MN survival assays. It would be interesting to determine whether those “pathological marks” are selectively present in the most vulnerable MN subpopulations in SMA patients.

While the corrected-SCO neural and MN longitudinal gene expression profiles changed in comparison to the SMA SCOs, trending towards healthy control values, this rescue effect was not complete and varied across the genes studied. Of note, the pan-neuronal marker SMI-32 profile was almost identical between the corrected and diseased SCOs, plateauing at an early stage although a gradual increase was expected, and detected in the healthy SCOs, if neurons continue to differentiate and mature. Also remarkable was the pronounced drop in the levels of the MNs markers ISL1 and CHAT, similar to the SMA SCOs. This observation raises the troubling possibility that not all neurological alterations in SMA patients might be solved by raising SMN protein, regardless of how early the SMN-increasing treatment is provided. The partial rescue effect observed in the corrected isogenic SCOs over the pathological newly identified SMA phenotypes arouses interesting hypotheses. First, it is possible that for all three SMA lines studied other factors, independent of SMN, play a role in the disease pathogenesis. It is possible that other genes, localized within the highly complex 500 kbp region were *SMN* genes sit also carry mutations, or that other genetic alterations are present in these lines contributing to the disease. This possibility is unlikely, however, only by comparing SMA lines to their isogenic controls, and not with different healthy lines, true disease contributors could be discovered. Alternatively, potential epigenetic alterations in the SMA lines caused by SMN deficiency, not fully erased during the fibroblasts reprogramming, are not corrected by replenishing the protein and are exerting important functions in the development of SMA pathology. This possibility is supported by three previous studies. Sabra et al found that SMN is a chromatin-binding protein that specifically interacts with methylated histone H3K79, a gene expression- and splicing-associated histone modification (Sabra et al., 2013). Further, a genome-wide methylation report on peripheral whole blood leukocytes from SMA and control individuals showed numerous CpG sites close to genes differentially methylated between patients and controls (Zheleznyakova et al., 2013). More recently, several genes involved in multiple stages of MN differentiation, also showed an increased methylation profile in SMA hiPSC-derived MN cultures compared to healthy controls, including *PAX6, OLIG2, HB9, ISL1* and *CHAT* (Maretina et al., 2022). The expression of these genes was not measured in this study; however, it is likely that this increased methylation alters their expression and therefore the specification, differentiation and/or maturation of the MNs. Other studies (Sanchez et al., 2013; Lee et al., 2022) have shown dysregulations in the abundance of several methyltransferases in SMA, that we have also detected in a MS study on HB9-purified MNs that we have performed (unpublished). Epigenetic modifications in SMA patients, SMN-dependent or independent, could be reason for the different disease phenotypes that patients carrying the same *SMN2* copy number present and for the variable response of SMA patients to treatments. Similarly, distinct epigenetics could also explain the partial restoration of the observed pathological phenotypes in our isogenic SCOs when compared to the parental SMA. Longitudinal scRNAseq/scATACseq studies together with complementary omics on these SMA and isogenic-control SCOs will be critical to unravel the molecular signatures responsible for selective MN vulnerability in SMA.

## Tables

**Table 1.** Primers used for PCR-amplification of SMN loci followed by Sanger sequencing.

**Table 2:** Primers used for qPCR mRNA expression quantification for hiPSC Trilineage differentiation assay (genes recommended by STEMdiff Trilineage Differentiation Kit) and for neural differentiation of SCOs.

## Materials and Methods

### Study approval

The healthy BJ siPSC line and the SMA hiPSCs were kindly provided by Lee L. Rubin (Harvard University) through an MTA. Subsequent use of the hiPSC lines in the Rodriguez-Muela lab at DZNE was approved by the Ethics Commission at the Technische Universität, Dresden (SR-EK 80022020). The information regarding both the SMA hiPSC lines and the healthy control line is enclosed in Rodriguez-Muela et al 2017 (Rodriguez-Muela et al., 2017).

### Generation of the SMN:Clover vector

Briefly, one plasmid (px458, Addgene Plasmid #48138) expressing sgRNA as well as Cas9 was used to introduce double-strand-breaks near Exon 7 of *SMN2* locus. A second plasmid, pTG-HR-SMN:Clover, a modified version of the commercially available plasmid HR120-PA1 (Systembio) served as targeting vector carrying *SMN1* exon 7 (C in position 6) as well as the Clover fluorophore and was used as template for homologous recombination to repair the double strand break post cleavage by Cas9. To generate the pTG-Cr-SMN, we designed a CRISPR guide (gRNA) with an estimated cleavage site right before the stop codon of exon 7 *SMN* loci. The following single-stranded (ss) oligos (IDT) were annealed and cloned into the px458 plasmid (Addgene # 48138) using BbsI and T7 DNA ligase in a one-step digestion-ligation reaction to produce the CRISPR gRNA.

gRNA 1 FOR for SMN-Clover: **CACC**TGCTCACATTCCTTAAATTA

gRNA 1 REV for SMN-Clover: **AAAC**TAATTTAAGGAATGTGAGCA

The correct insertion of the gRNA into the px458 plasmid, resulting in pTG-**Cr**-SMN, was confirmed by Sanger Sequencing (Microsynths). To generate pTG-HR-SMN:Clover, the commercially available vector HR120-PA1 (Systembio) was used as backbone. First, HR120-PA1 was digested with EcoR1 and NRU1 to remove copGFP, WPRE and PolyA. One gBlock (IDT) was then used to insert Clover coding sequence and to restore WPRE and PolyA as well as EcoR1 and NRUI restriction sites. Following another digest with EcoR1, a gBlock containing the last 372bps of intron 6 (of *SMN2*) and *SMN1* exon 7 without its stop codon - used as the 5’ homology arm (HA) and was cloned into the vector via Gibson assembly. Finally, the vector was digested with BamHI to introduce a second gBlock containing the 3’ HA -consisting of 420bp of *SMN2* intron 7 (Table 2). The successful cloning of both HAs as well as the correct sequences of the inserted gBlocks were confirmed via Sanger sequencing. The final vector also contains the double selection cassette, mRuby-T2A-puromycin, flanked by loxP sites, to enable preselection for successful targeted cells.

### Generation of the isogenic corrected hiPSCs from SMA lines

For each of the hiPSC lines 1*10^6 cells were dissociated using Accutase and resuspended in 100 μl P3 nucleofection solution. 2.5 μg of the pTG-Cr-SMN plasmid and 5 μg of targeting vector pTG-HR-SMN:Clover were added. Nucleofection was performed using the 4-D nucleofector system (AMAXA) and the P3 Primary Cell 4D-Nucleofector Kit (Lonza, V4XP-3024) following manufactures’ instructions (program CB-150). hiPSCs were transferred to matrigel (Corning, 354234)-coated dishes containing (Stem Cell Technologies; 85850) with 4 μM ROCK inhibitor (Hölzel Diagnostik; S1049-50) to improve survival. ROCK inhibitor was removed 24 h post nucleofection and media was change from there on every other day. From d2-d9 1 µg/ml puromycin (Life Technologies, A1113802) was added to select for successful targeted cells. At day 12, the cells were nucleofected with a CRE-GFP plasmid to excise mRuby-T2A-Puromycin. 36 hours post nucleofection GFP+:mRuby-cells were FACsorted and plated on matrigel-coated 10cm dishes at different clonal densities. After 10 days, 24 colonies for each line were picked, expanded and sequenced. To confirm successfully targeted *SMN2* loci, as well as presence of untargeted *SMN2* the PCR strategy shown in Figure S1B-C was used. Briefly, to check for untargeted *SMN2* copies forward primer upstream of the 5’ HA in combination with reverse primer in intron 7 was used. To check for targeted SMN:Clover loci the same forward primer was used, while the reverse primer was designed to bind within Clover sequence therefore only giving a product if Clover had been fused to exon 7 of the corrected *SMN2* locus. The primers used are contained in Table 1. At least four clones per line were successfully sequenced, having at least one *SMN2* copy targeted (mutation corrected and Clover added in frame) and one copy that remained untargeted. Two of the successfully targeted clones per SMA line were used in this study.

### Validation of correct genome editing by PCR followed by Sanger Sequencing

hiPSCs were collected, washed and spun down at 135g for 5 min. Cells pellets were lysed and gDNA extracted using DirectPCR lysis reagent CELL (Viagen, 301-C) and Proteinase K (Thermo Fisher Scientific, EO0492). PCRs to confirm targeted and untargeted *SMN2* loci were performed for each of the cell lines with primer pairs shown in table 1 using High-Fidelity 2X PCR Master Mix (NEB, M0541L) and Thermal Cycler C1000 Touch (Bio-RAD). Using the same forward primer binding upstream of the 5’ homology arm, two different PCRs were performed. First, to confirm successful targeting and therefore conversion of *SMN2* into *SMN1*, a reverse primer binding within the Clover sequence was used. Second, to confirm the presence of untargeted alleles, a reverse primer binding in intron 7 was used. Due to the distance from the forward primer, this second PCR only worked if no Clover had been inserted. To confirm successful PCRs, samples were run in 1.5% (Sigma-Aldrich, A9539) gels stained with 0.01 % RedSafe (INtron Biotechnology, 21141) using Perfect Blue Gel System (Peqlab). PCR products were analyzed by Sanger sequencing (Microsynths) for final confirmation of targeted and untargeted *SMN2* alleles for each cell line.

### Human iPSC protein turnover and proliferation assays

To determine total SMN (antibody detected), SMN:Clover and Gemin2 protein turnover, 4.000 hiPSCs per line were plated in matrigel-coated 96W plates and 2 days later and incubated with 0.5 µg/ml of cycloheximide (CHX, Sigma Aldrich, C1988) for the indicated time followed by fixation, permeabilization and immunostained. Imaging and image analysis was performed as described below. Proliferation rate of the different hiPSC lines was determined using live imaging and automated high throughput analysis. Cultures of hiPSCs (at ∼60-80% confluency) were dissociated to single cell suspensions using accutase (Corning, 25-058-Cl), washed in 1x Dulbecos PBS-Ca/-Mg (Thermofisher Scientific; 14190169), spun at room temperature (RT) 135g for 5 minutes, and plated to Black F-Bottom Greiner µClear p96 well plates (Greiner, 655090) at a density of 4.000 cells/well. The plating medium consisted of mTeSR1, 5x mTeSR1 Supplement, 5% PenStep (Thermofisher Scientific, 1510-122), and 10µM Y-27632 2 HCL (ROCKi). hiPSCs were maintained with daily media changes with mTeSR1. 2 days post plating (D2) cells were stained with 500nM of SiR-DNA (Spyrochrome, SC007) dye for 1-1.5hrs before removal and 2x 1x PBS washes. After staining cells each well was imaged in the CO2 and temperature controlled Operetta CLS (5% CO_2_ and 37°C) with 20X water immersion objective (NA 1.0; Plan Apochromat) and Alexa647 emission filter. On subsequent imaging days, cells were stained with 500nM SiR-DNA for 30min to refresh effluxed dye and when cells reach 95-100% confluency the experiment was ended. To quantify iPSC proliferation, the Perkin Elmer software Harmony v4.0 was used to perform automated analysis of colony area µm^2^ at each time point. This was achieved by first applying an image filter (mean smoothing filter or sliding parabola) to clarify whole colonies in the Alexa647 channel. SiR-DNA+ Image Regions were found on these filtered images, and only living regions selected. The resulting selected areas were used as the final measurement.

### hiPSC differentiation into the 3-germ layers

hiPSCs were plated in matrigel-coated 24-well plates and cultured in STEMdiff Trilineage Ectoderm, Mesoderm or Endoderm differentiation medium according to manufacturer’s instructions (STEMCELL, 5230). To measure gene expression, cells were washed 2x PBS and lysed in TRIzol. RNA isolation and mRNA expression was determined as described in the “RNA-Isolation, cDNA-synthesis and qRT-PCR” section.

### RNA-Isolation, cDNA-synthesis and qRT-PCR

hiPSCs or SCOs were washed 2x PBS. Total RNA was extracted with the TRIzol reagent (Life Technologies, 15596026), and the concentration was measured with the NanoDrop TM 1000 Spectrophotometer (Thermo Scientific). 0.5-1 μg RNA was subjected DNase I digestion using (Thermofisher Scientific, EN0521) according to the manufacturer’s instructions. RNA was reverse transcribed with High-Capacity cDNA Reverse Transcription Kit (Applied Biosystems, 4368814). qRT-PCR was performed with GoTaq qPCR Master Mix (Promega, A6002) and a Quantstudio5 Real-Time PCR Detection System (ThermoFisher). mRNA expression levels were normalized to the expression of the human housekeeping gene h18S and the cycle numbers plotted (hiPSC Trilineage Differentiation) or relative values were determined with the comparative ddCT method (SCO developmental gene expression). Ultimately, the gene expression was normalized against the Control group of each experiment. Primers used are depicted in Table 2. 3 independent hiPSC Trilineage Differentiation experiments were analyzed. 4-8 SCOs (depending on the timepoint, 4 for d28 and d38, 6 for d18 and 8 SCOs for d8) per experiment were pooled together for RNA isolation, from 4 independent experiments.

### Human spinal MN differentiation

hiPSCs were grown on cell culture treated dishes, coated with Matrigel, maintained with mTeSR1 and split using ReLeSR (StemCell, 100-0484). For initiation of the MN differentiation protocol, hiPSCs were dissociated and cultured in mTeSR as embryoid bodies (EBs) for 3 days in ultra-low attachment dishes (ULA) (Corning, 3262). For the first 24 hours mTeSR was supplemented with 10µM Y-27632 (RHOK inhibitor) and 10ng/mL FGF-2 (Millipore, 32160702). 3 days later the media was changed to neuronal induction media (NIM) (day 0), containing 50% Advanced DMEM/F12 (ThermoFisher, 12634028), 50% Neurobasal media (ThermoFisher, 21103049), 1% Penicillin-Streptomycin (LifeTechnologies, 15140122), 1% GlutaMAX (ThermoFisher, 35050087), 0.1mM 2-ß-mercaptoethanol (LifeTechnologies, 21985023), 0.5x B27 (ThermoFisher, 17504044) 0.5x N2 (ThermoFisher, 17502048), and 20µM ascorbic acid (SigmaAldrich, A4403). From day 0 to day 4 NIM media was supplemented with 10µM SB 431542 (BioTechne, 1614/10) and 100nM LDN 193189 (Hölzel, M1873) to induce neural differentiation. From day 3 to day 15 1µM retinoic acid (SigmaAldrich, R2625) and 1µM Sonic Hedgehog Signaling Agonist (Millipore, 566660) were added. From day 7, 5ng/mL BDNF (R&D, 248-BD-025/CF), from day 9 10µM DAPT (Tocris, 2634/10) and from day 11 5ng/mL GDNF (R&D, 212GD0/50CF) and 2µM cytosine arabinoside (AraC, SigmaAldrich, C1768) were added. On day 15-17 the EBs were dissociated with papain/DNase solution (Worthington, LK003178, LK003172) and plated on 50µg/mL poly-D-lysine (SigmaAldrich, A-003-E), 3µg/mL laminin (ThermoFisher, 23017015) coated plates. Media for culturing dissociated neurons was Neurobasal containing 1% Penicillin-Streptomycin, 1% GlutaMAX, 1x non-essential amino acids (LifeTechnologies, 11140050), 0.5x B27, 0.5x N2, 20µM ascorbic acid, 25µM 2-ß-mercaptoethanol, 2µM AraC, 10ng/mL BDNF and 10ng/mL GDNF. A full media change was performed 2 days after plating, then half media changes were performed every second to third day.

### Human spinal MN treatment, survival assays and image analysis

When indicated, MNs were treated with 1 µM MLN4924 (Active Biochem) (to stabilize SMN protein), 0.3µg/ml CHX (to prevent protein synthesis and therefore detect protein degradation) or DMSO control 4 days after being plated for 3 days. Cells were then fixed for 15 min in 4% PFA, permeabilized for 30 minutes (0.25% TritonX (Sigma, X100) and 5% NGS (Cell Signaling, 5424S), immunostained with the indicated primary antibodies for 2h at RT and secondary antibodies for 1h at RT. Images were captured using an automated Operetta CLS microscope (PerkinElmer) with water immersion-40X magnification. Subsequent image quantification was performed using the Columbus Analysis System. MNs were identified by ISL1 fluorescence in the nucleus and nuclei were identified using Hoechst. Total SMN or SMN:Clover fluorescence intensity was determined as previously described (Rodriguez-Muela et al., 2017). Briefly, after cell-identification a constant cytoplasmic region (circle) around the nucleus was defined and, by using a fully automated imager and associated software, the intensities in the SMN-immunostained and SMN:Clover channels of all the pixels in that cell region were added up giving the “total intensity” in arbitrary units. That number was then divided by the number of pixels in that region, resulting in the “mean intensity” (of a pixel in the cell) in a way that is independent of the cell size. The mean intensity per cell was averaged across 40 random fields per well and 3-5 wells per condition, containing in total hundreds to thousands of MNs in each experiment. For each experiment, three wells with no primary antibody or non-Clover cells were used to determine background fluorescence intensity. To assess survival on live MN cultures using SiR-DNA (Figure 3A-E), after EB dissociation, neurons were plated in poly-D-lysin/laminin-coated 96 well plates (Corning, 4680) at a density of 8*10^4^ cells/well. 3 wells per cell line were stained using the live nuclear dye SiR-DNA at 125nM for 1.5h on day 2 after plating and washed with 1X PBS. Staining of the cells was repeated every 3 days with 62.5nM SiR-DNA dye for 1h. To follow survival of the neuronal cultures, stained wells were imaged every 2 days using the Operetta CLS microscope with water immersion-40X objective. Image quantification was performed using the Columbus Analysis System. A size and morphology threshold was used to identify living cells and to eliminate apoptotic nuclei from quantification. To validate the increased survival of MNs derived from the most severe SMA line (Figure 3F-G), MNs were plated in poly-D-lysin/laminin-coated plates that were fixed 2 and 12 days later. The number of ISL1+ MNs was quantified at both time-points and the percentage of surviving MNs over that timeframe for each line graphed.

### Generation of SCOs, live imaging and longitudinal size quantification

hiPSCs colonies from the 10 lines (BJ WT and the 3 isogenic trios) were accutased and 4.000 cells per well seeded into ULA 96-well plates and cultured in mTeSR for 5 days (ROCKi was added for the first 24 hours). The formed stem cell aggregate were changed to neural induction medium (NIM) and 3 days later embedded in 15µl matrigel drops, transferred to ULA 10 cm dishes and kept in an orbital shaker, where the spinal cord patterning started. Dual SMAD inhibition (10µM SB431542 + 100 nM LDN193189) was maintained for 6 days (day 0 to day 6). On day 4 the caudalizing agent RA (1µM) and the ventralizing SAG (1µM) were added for 12 days (day 4 to day 16). The notch response inhibitor DAPT (2.5µM) was added to the culture on day 10 to enhance neural differentiation (from day 10 to day 16). Neurotrophic factors were added subsequently and until the end of the culture (10nM BDNF from day 12 and 10nM GDNF from day18). To image stem cell aggregate formation and sphere growth, the 96w ULA plates were imaged two days after being seeded (day -6) in a Operetta CLS microscope, under temperature and C02 control, using a 10x air objective and imaging 10 focal planes 20µm apart, every day for 6 days. Sphere size quantification was performed blindly on images with maximum projection of all Z-stack sections using ImageJ software (NIH). 24-36 spheres per hiPSC line, day and experiment were measured (3-5 experiments per line).

### Cryosectioning, immunofluorescence staining, imaging and image analysis of SCO sections

After overnight fixation in 4% PFA, SCOs were washed o/n in 1x PBS and cryopreserved gradually in sucrose before being embedded (TissueTek OCT, Fisher Scientific, 10690461) and flash-frozen on dry-ice. The cryoblocks were sectioned at 15µm thickness using the Cryostat Leica CM3050S (Leica). Sections were collected onto SuperFrost Plus Slides (Thermo Scientific) and kept at -80°C. For immunofluorescence staining of SCO sections, the slides were post-fixed in 4% PFA for 10 min, washed and blocked for 1h with Blocking Buffer (BB) (0.25% TritonX, 5% NGS, 0.1% Tween20 in 1x PBS) at RT in a humid chamber. Primary antibodies diluted in BB were added for 2h at RT. Slides were washed and secondary antibodies added for 1h at RT. After several washes, Hoechst33342 (Life Technologies, H3570) was added and the slides mounted on coverslips with Vectashield Antifade Mounting Medium (Biozol Diagnostica, VEC-H-1000). Imaging was carried out using a Zeiss Confocal Spinning Disc microscope using multi-slide holders and automatic scan of tile regions, with a 20x PlanApoChromat Air Objective (NA=0.8, DICII). Z-Stack scans of 15µm thickness with an interval of 1µm were recorded for each organoid section. The same imaging parameters were used across all SCOs derived from all hiPSC lines for each of the antibodies used. Maximum intensity projections of z-Stacks were made and stitched with 10% tile overlap by Hoechst33342 as reference channel. Automatic quantification of the number of cells positive for the marker of interest was carried out in Arivis Vision4D 3.3.0 (Zeiss). Briefly, CZI.files were converted into arivisSIS format with no compression, using Zen 3.2. The quantification pipeline in Arivis Vision4D 3.3.0 included: 1) a fluorescence threshold to exclude background (based on 2ry-only stained SCOs), 2) global enhancement “Simple Sharpening Filter”, 3) BlobFinder-nuclear segmentation tool for Hoechst33342-Channel and 4) segment operation “Feature-Filter” on surface-area (µm²) to remove small artifacts. Quantification was carried out using “Batch Analysis” for all SCOs. 3-4 SCOs per experiment were analyzed from 3-6 independent experiments.

### Co-Immunoprecipitation Assay

hiPSCs were lysed in lysis buffer (25mM Tris HCl pH 7.4, 62.5mM NaCl, 2% NP-40 (Thermofisher Scientific 85124), 1mM EDTA (Thermofisher Scientific, 15575020), 100x HALT protease inhibitor cocktail (Thermofisher Scientific, 1861278), and 0.03 U/µl DNase in 1x PBS). Cells were lifted from the plate using ReLeSR (Stem Cell Technologies; 100-0484) and hiPSCs were spun for 5min at RT and 135g. Samples were placed on ice and mechanically lysed in pre-chilled lysis buffer using pre-chilled pipette tips and passing through pre-chilled syringes 10 times. Lysate was incubated on ice for 20min before spinning down for 10min at 13.523g and 4°C. 1 mg of protein was used in each reaction, and total lysate was taken from each sample prior to equalizing the reaction volume to 500µL with Co-IP wash buffer (lysis buffer without Protease Inhibitor or DNase). 15µL of GFP-Trap DynaBeads (Chromotek, gtd-20), 2µg anti-SMN antibody or 0.1µg anti-IgG mouse were added to each reaction. Reactions were incubated overnight at 4°C with rotation, and then 15µL of Pierce Protein A/G Magnetic Beads (Thermofisher Scientific, 88803) were added to the anti-SMN and anti-IgG reactions and incubated for 2 hours at 4°C with rotation. Following bead incubation, beads were separated from supernatant and washed 3x with pre-chilled wash buffer and 3x with pre-chilled 1x PBS for 10min each at 4°C with rotation. Following washes, proteins were eluted from beads with 2x Laemli Buffer (Bio-RAD, 1610737) with β-mercaptoethanol and boiled at 95°C for 15minutes along with total lysate samples. Samples were run on western blot following the described procedure below and proteins were probed for the indicated antibodies.

### Western Blot

Cells lysed in RIPA buffer (Thermo Fisher Scientific, 89900) with complete protease and phosphatase inhibitors as described above. Western blots were performed using AnykD Criterion TGX Precast Midi Protein Gels (Bio-RAD, 5671124, 5671123). Gels were run in 1x Tris-Glycine SDS Running Buffer (Thermofisher Scientific, LC26755) using 60V for ∼30min and 100-110V for ∼2hours. After running, the gel was equilibrated in 1x Tris/Glycine Transfer Buffer (Bio-RAD, 1610734) for ∼5min. Proteins were transferred from gel to pre-made TransBlot Turbo-Transfer Pack (Bio-RAD, 1704157) in the semi-dry Trans-Blot Turbo System (Bio-RAD, 1704150). Once transferred, the membrane was rinsed with diH_2_O and stained with Ponceau to confirm even loading. Membranes were then blocked with 5% nonfat dried milk powder (PanReac Applichem ITW Reagents, A0830) in 1x TBST (ChemCruz, sc362311) for 1 hour at RT and agitation. The desired antibody was then added and incubated at 4°C overnight; the next day the membrane was washed 3x with 1x TBST for 10min each, probed for the secondary antibody diluted in 5% milk at RT with shaking, and washed again. The signal was developed using western blot substrate (SupraSignal West PICO PLUS Chemiluminescent Substrate; Thermofisher Scientific, 34580), X-Ray Films (FUJI, 4741019289), and the Cawomat 2000 IR X-Ray developer. Densitometric analysis was performed on scanned autoradiographs using the Quantity One software (Bio-RAD).

### Primary antibodies used

anti-SMN (for co-IP) (Novus Biologics, NB100-193); anti-IgG mouse (Santa Cruz, sc2025); anti-SMN (for immunofluorescence) (BD Biosciences, 610646); anti-Gemin2 (Abcam, ab150383); anti-SOX2 (Invitrogen, 14-9811-82); anti-NESTIN (Millipore, MAB353); anti-ISL1 (Abcam ab109517); anti-NKX6.1 (Developmental Studies Hybridoma Bank, F55A10); anti-MAP2, NovusBiologics NBP2-25156); anti-β-Actin (Cell Signaling, 8H10D10), anti-β-Tubulin (Cell Signaling, 2146).

### Statistical Analysis

Statistical significance was determined using GraphPad Prism 9.3.1 (Graphpad Software, Inc.). To test the Gaussian distribution of residuals Shapiro-Wilk test was performed. To test equal distribution of standard deviations (SD) Bartlett’s test was performed. If no Gaussian distribution of the residuals, nonparametric Kruskal-Wallis test was performed. If no equal SD among groups, mixed Brown-Forsythe and Welch ANOVA tests were performed. One-way ANOVA was performed for datasets composed of only one variable (e.g. genotype) and Two-way ANOVA was chosen for datasets composed of two variables (e.g. genotype and time during differentiation). A confidence interval of 95% was used for all comparisons. Graphs indicate mean+SEM.

## Acknowledgments

We would like to thank Patricia Vazquez and Federico Calegari for constructive comments and feedback. We thank Lee Rubin (Harvard University) for providing the MTA enabling the work with the BJ WT and SMA parental hiPSCs. We thank Eric Geertsma (MPI-CBG), Aliona Bogdanova (Protein Biochemistry Facility, MPI-CBG) and Marc Gentzel (Mass Spectrometry Facility, CMCB) for insightful discussions on SMN:Clover protein stability. We thank Silke Winkler (Sequencing Facility, MPI-CBG and DcGC) for helpful advice on PCR amplification and sequencing of *SMN* loci. We thank Eugster Oegema (Organoid and Stem Cell Facility, MPI-CBG) for her assistance with hiPSC karyotyping. We thank Silke White (DZNE Imaging Facility) and Ellen Geibelt (Light Microscopy Facility, CMCB) for their support and training on multiple microscopes. We would like to acknowledge the work of Katharina Nolte, Irmgard Hölker and Anixa-Muinos-Bühl in determining the *SMN1* and *SMN2* copies in all cell lines. This work was supported by the European Research Council (ERC-StG 802182), Helmholtz (AMPro, ZT-0026), German Society for Muscle Diseases (DGM, Bu4/1), Funding Programs for DZNE-Helmholtz, TU Dresden CRTD and MPI-CBG to NRM. This work was also supported by the German Research Foundation [Wi 945/17-1 and CRC1451 (project-ID 431549029 – A01), and European Union’s Horizon 2020 research and innovation program under the Marie Skłodowska-Curie grant agreement No 956185 (SMABEYOND) and Center for Molecular Medicine Cologne (project No C18) to BW. The authors declare no competing financial interests.

## Author Contributions

TG and NRM designed the experiments. TG, IR, JT, FB, ZD and NRM performed the experiments. AD provided technical assistance. BW and JB determined the SMN1 and SMN2 copy number. TG, IR, JT, FB, ZD and NRM analyzed the data. TG and NRM wrote the manuscript. BW and NRM obtained funding. All authors provided editorial comments.

