## Supplemental Figures 1-5 for "An isogenic human iPSC model unravels neurodevelopmental abnormalities in SMA"

**Supplemental Figure 1. Generation of isogenic control hiPSCs from various severities of SMA human iPSC lines** (associated to Figure 1). (**A**) Scheme of pTG-HR-SMN:Clover targeting vector used for knock-in mutagenesis to generate the isogenic hiPSC lines from the SMA parental lines. (**B**) In silico design to detect a non-targeted *SMN2* copy in a parental SMA line and chromatograms from the targeted lines showing a non-edited *SMN2* copy. *SMN2* locus is indicated by the blue bar. Exon7 of *SMN2* is labeled by a yellow bar. The primers used for PCR amplification covering the regions are indicated by the green triangles and the sizes of the resulting gDNA fragments are indicated below. (**C**) Electrophoresis gel showing the PCR products corresponding to primer pair shown in (B) for the healthy control untargeted BJ WT hiPSC line as well as targeted BJ WT, all 3 untargeted SMA parental lines as well as two targeted *SMN1*:Clover clones per line. The gel shows that for each line at least one *SMN2* locus remained unedited. (**D**) In silico design to detect a successfully targeted *SMN2* copy in a parental SMA line and chromatograms from the targeted lines showing an edited *SMN2* copy. *SMN2* locus is indicated by the blue bar. Exon7 of *SMN2* is labeled by a yellow bar. The 5' and 3' gBlocks used for the homologous recombination are indicated by pink bars, the Clover sequence (which is part of the targeting vector) is shown in green. The primers used for PCR amplification covering the region are indicated by the green triangles and the size of the resulting gDNA fragments are indicated below. (**E**) Electrophoresis gel showing that for each of the clones at least one *SMN2* locus has been successfully targeted and converted to *SMN1*:Clover. (**F**) Representative images from the hiPSCs used in the study fixed and immunostained against the pluripotency markers OCT4 (turquois), SOX2 (magenta), Tra-1-60 (orange) and Nanog (green). Nuclei were identified with Hoechst. Scale bar, 50 µm.

**Supplemental Figure 2** (associated to Figure 1 and 2). **The Clover reporter tagging SMN C-terminus does not alter the biology of the protein.** (**A**) Quantification of the fluorescence intensity of nuclear gems endogenously identified by SMN:Clover or immunostained using an anti-SMN antibody (**B**) in MN cultures generated from the healthy BJ WT line and two BJ SMN:Clover clones ("BJ #16" and "BJ #23"), 4 and 8 days after being plated (**$p<0.01$, ****$p<0.001$ One-way ANOVA followed by Tukey's multiple comparison test, n=3). (**C**) Quantification of the number of SMN:Clover+ (white bars) or SMN-antibody detected (grey bars) nuclear gems per ISL1+ MNs in MN cultures generated from two BJ SMN:Clover edited clones 4 days after being plated with 1 µM MLN4924 or 0.3µg/ml CHX (*$p<0.05$ One-way ANOVA followed by Tukey's multiple comparison test, n=4). Representative images are shown in (**D**) (SMN, red; ISL1, cyan; Hoechst, blue; endogenous SMN:Clover, green).  Scale bar, 10 µm. Quantification of the SMN fluorescence intensity detected by antibody labeling, in hiPSCs treated with CHX for the indicated times (in hours) related to the DMSO-treated cells (Ct) to determine total SMN protein (Clover-tagged and untagged) half-life in the healthy control BJ (**E**), SMA type III (**F**), type II (**G**) and type I (**H**). Quantification of the number of hiPSCs treated with CHX for the indicated times (in hours) related to the DMSO- treated cells (Ct) to determine cell toxicity caused by the treatment in the healthy control BJ (**I**) SMA 39C-III and its corrected clones (**J**), 51N-II and its corrected clones (**K**) and 38D-I and its corrected clones (**L**).

**Supplemental Figure 3** (associated to Figure 3). **The corrected isogenic SMA hiPSCs differentiate into the three-germ layer similarly to the SMA parental and healthy control hiPSC lines. (A)** Representative images showing SiR-DNA labeling of type III and II isogenic iPSC trios 2, 4 and 6 days after plated in Matrigel-coated 96w plates. Scale bar, 1 mm. (**B**) Exemplification of the image processing steps done by the Harmony v4.0 software script to detect the SiR-DNA labeled hiPSCs from all imaged wells in each well, filter the images by applying a Sliding Parabola algorithm (2) or a Mean Smoothing algorithm (3) and then identify the area occupied by the whole Image Region (4). qPCR quantification of expression makers of ectodermal *SOX1* (**C**) and *PAX6* (**D**), endodermal *SOX17* (**E**) and *LEFTY1* (**F**) and mesodermal *NCAM* (**G**), *TBXT* (**H**) and *CXCR4* (**I**) markers from BJ WT hiPSCs cultured in STEMdiff™ Trilineage Differentiation Kit (STEMCELL). Graphics represent CT mRNA expression levels of the indicated genes subtracted to the housekeeping gene (18s) expression levels. The colored symbols in each graph represents the specific medium-cultured hiPSCs where the highest expression of that gene is expected (*$p<0.05$, **$p<0.01$, ***$p<0.005$, ****$p<0.0001$ One-way ANOVA followed by Tukey's multiple comparison test, n=4). mRNA expression qPCR quantification of ectodermal, endodermal and mesodermal marker genes from hiPSC SMA type III (**J**), type II (**K)** and type I (**L**) and their isogenic corrected clones cultured in STEMdiff™ Trilineage Differentiation Kit. Graphics represent CT mRNA expression levels of the indicated genes subtracted to the housekeeping gene (18s) expression levels (*$p<0.05$, **$p<0.01$, ***$p<0.005$ One-way ANOVA followed by Tukey's multiple comparison test, n=4).

**Supplemental Figure 4** (associated to Figure 4). **Similar percentage of ISL1+ and MAP2+ cells in the isogenic corrected 38D clones compared to the parental SMA lines 38D.** (**A**) Schematic representation of the protocol followed to generate hiPSC-derived spinal MNs. Representative images of BJ WT, SMA types III (**B**) and II (**C**) and both their respective isogenic corrected iPSC-derived MN cultures stained with SiR-DNA (red) after 2 and 10 days in culture. Bright field images of the corresponding neurons are also shown. Scale bar 20 µm. (**D**) Percentage of MAP2+ cells and ISL1+ MNs (**E**) in the MN cultures derived from the SMA 38D-I and both isogenic corrected hiPSC lines 2 days after being plated (One-way ANOVA test followed by Tukey's analysis, n=10).

**Supplemental Figure 5** (associated to Figure 5). **SMA hiPSCs are less efficient at self-assembling into spheres than the isogenic corrected clones or a healthy control line.** (**A**) Representative ULA 96w plate containing one stem cell aggregate per well 7 days after 4.000 51N-II SMA hiPSCs were seeded to illustrate the robustness of the culture. (**B**) Quantification of the area ($mm^2$) of SMA 39C-III and isogenic corrected hiPSC-derived spheres 1 day after the hiPSCs were seeded and until day 6. The violin plot exemplifies the size distribution of the individual spheres quantified for one representative experiment (****$p<0.0001$ Two-way ANOVA test followed by Tukey's analysis, n=30 spheres). (**C**) Time-course quantification of the BJ WT and SMA hiPSC-derived sphere size ($mm^2$) representing the average of at least 30 spheres imaged and quantified per line and per experiment (4-8 experiments) relative to the average sphere size at day 2 for each of the lines, (**D**) 39C-III parental and isogenic corrected clones sphere relative size growth, (**E**) 51N-II and isogenic corrected clones and (**F**) 38D-I and isogenic corrected clones (*p<0.05, **p<0.01 Two-way ANOVA test followed by Tukey's analysis; statistical significance between the BJ WT and the SMA lines -F- or the SMA and the corrected clones -G,H,I- is shown. # represents comparisons between BT WT and 39C-III; § between BT WT and 51N-II, n=3-6). (**G**) Representative images of the self-assembled spheres generated from BJ WT and the SMA hiPSCs 5 days after seeded. An increasing number of hiPSCs was seeded as a single-cell suspension in ULA 96w plates to determine efficiency of the hiPSCs to self-assemble depending on the starting cell number. Scale bar 400 µm. (**H**) Same spheres as shown in (J) 10 days later. (**I**) Representative images showing NESTIN expression (magenta) on day 8 SCOs from BJ WT, SMA 38D-I and corrected isogenic clones. Representative images showing MAP2 (magenta) (**J**) and ISL1 expression (white) (**K**) on day 28 SCOs from BJ WT, SMA 38D-I and corrected isogenic clones. Nuclei are labeled with Hoechst (blue) and the SCO border is indicated by a dotted line. Scale bar 100 µm. Quantification of the number of ISL1 (**L**) cells relative to the total number of cells per SCO in the BJ WT, SMA 38D-I and corrected isogenic clones. Values are shown relative to BJ WT (*p<0.05, **p<0.01, ***p<0.005 One-way ANOVA followed by Fisher's LSD multiple comparison test, n=3-6, 3-4 SCOs per experiment).

#### Supplemental Figure 1.

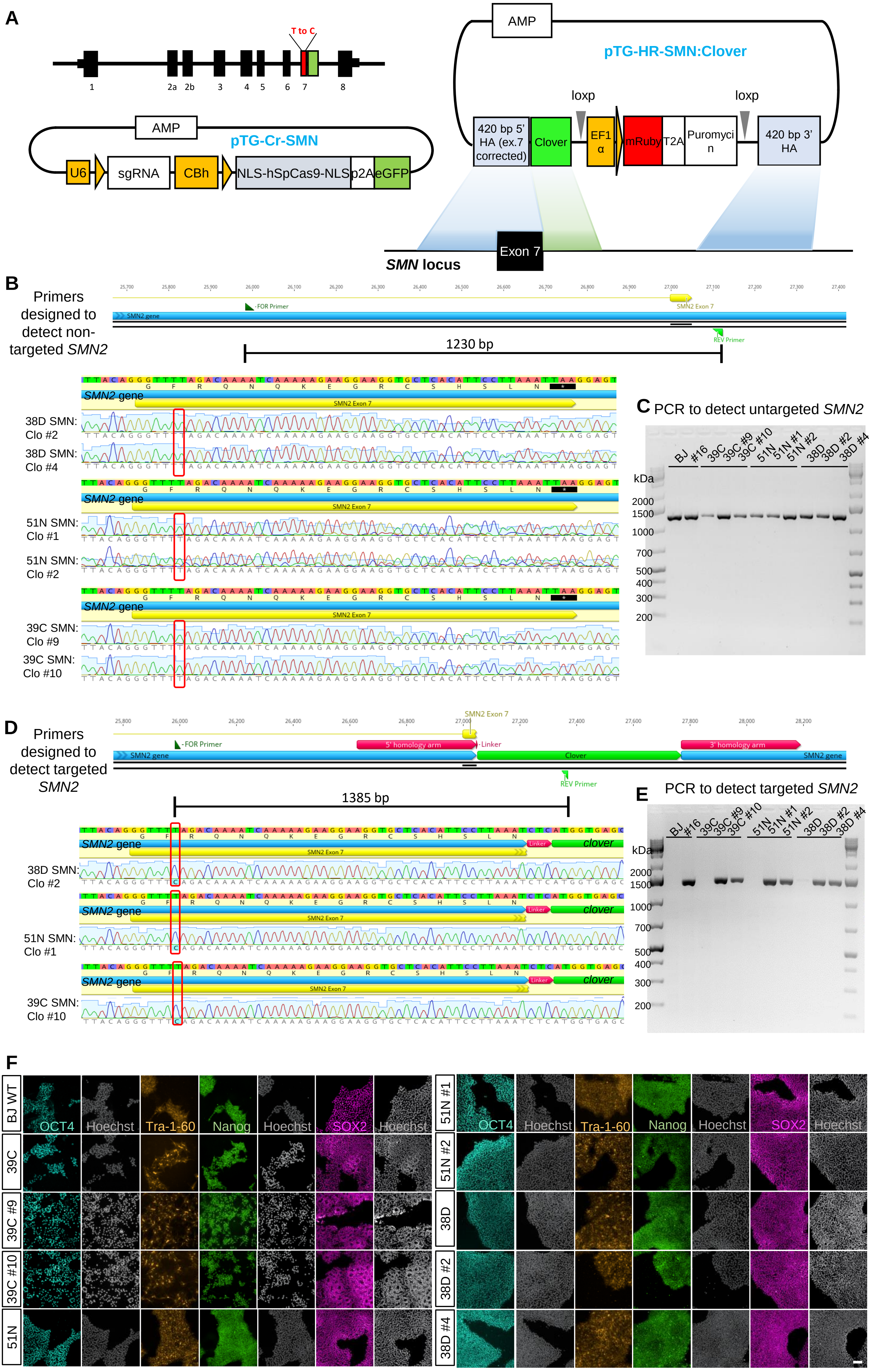

**Supplemental Figure 2.**

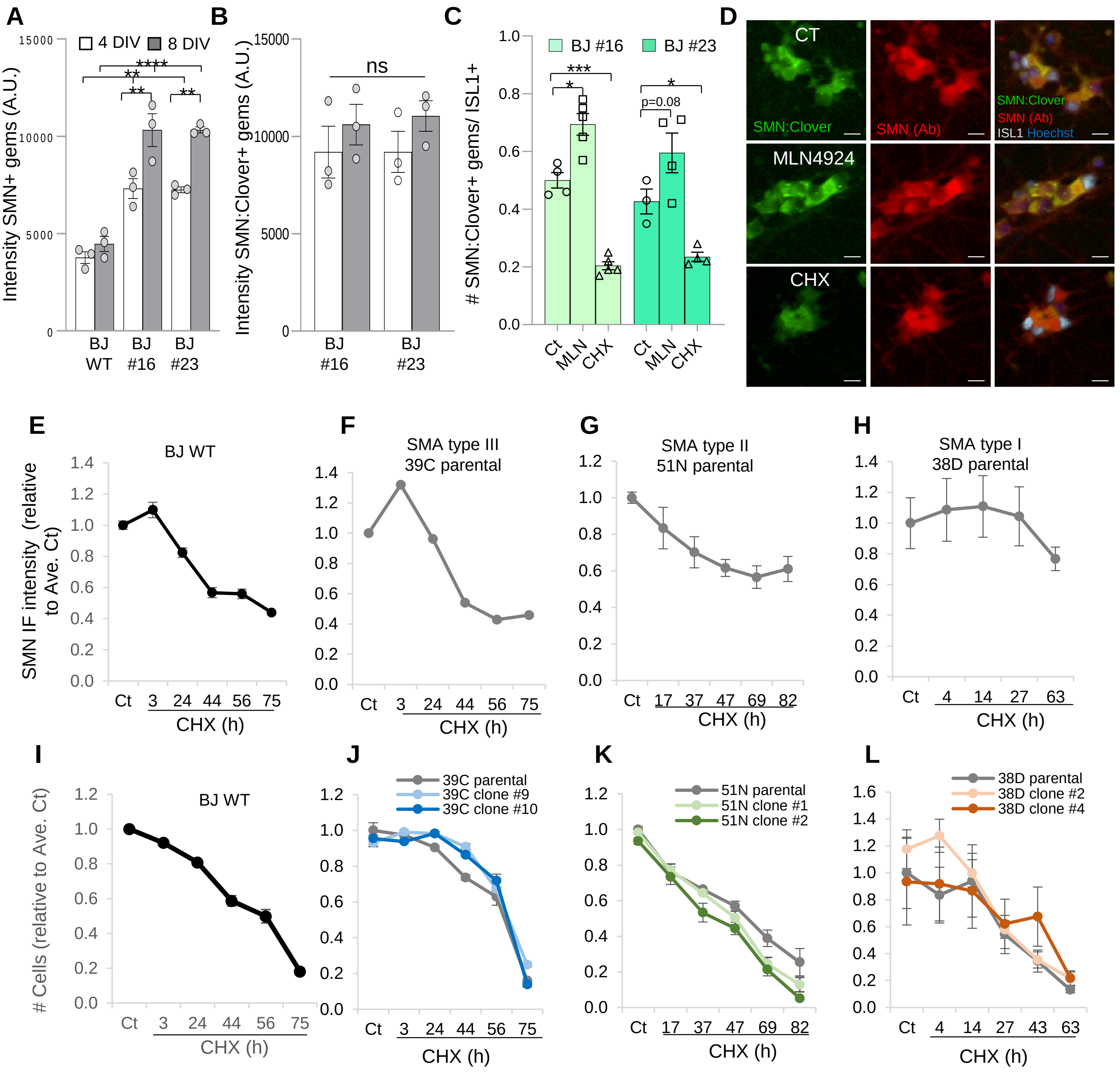

**Supplemental Figure 3.**

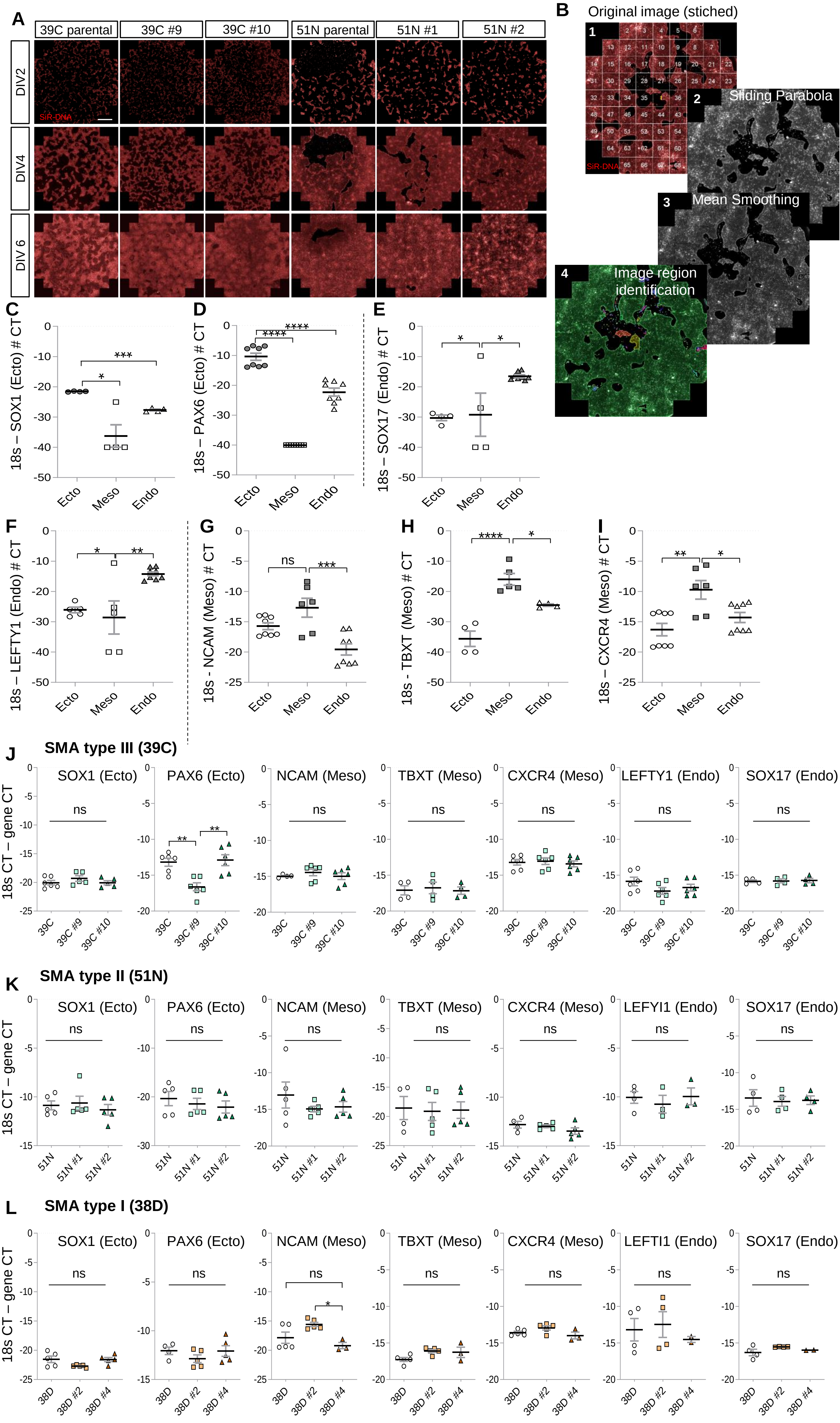

Supplemental Figure 4.

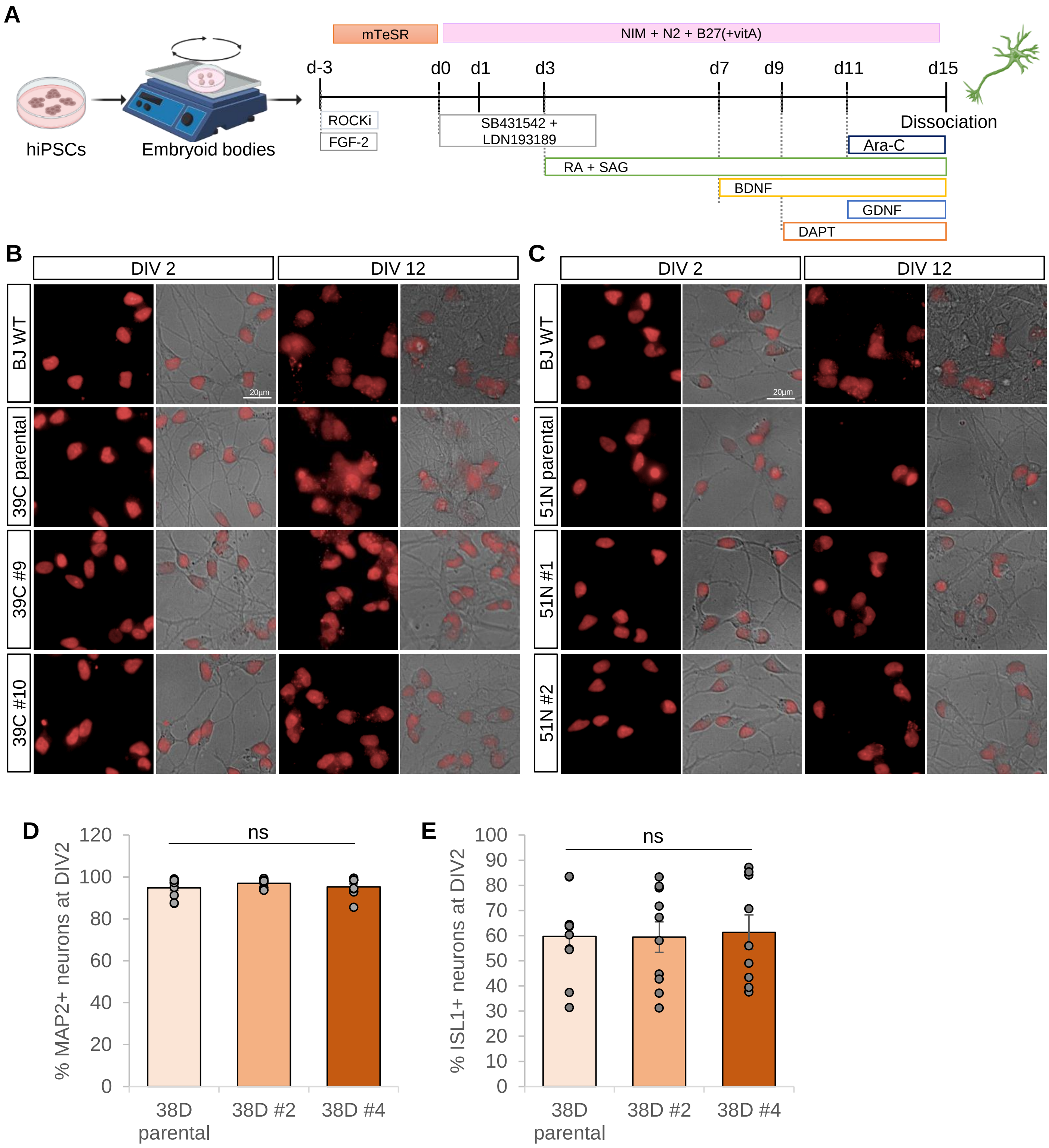

### Supplemental Figure 5.

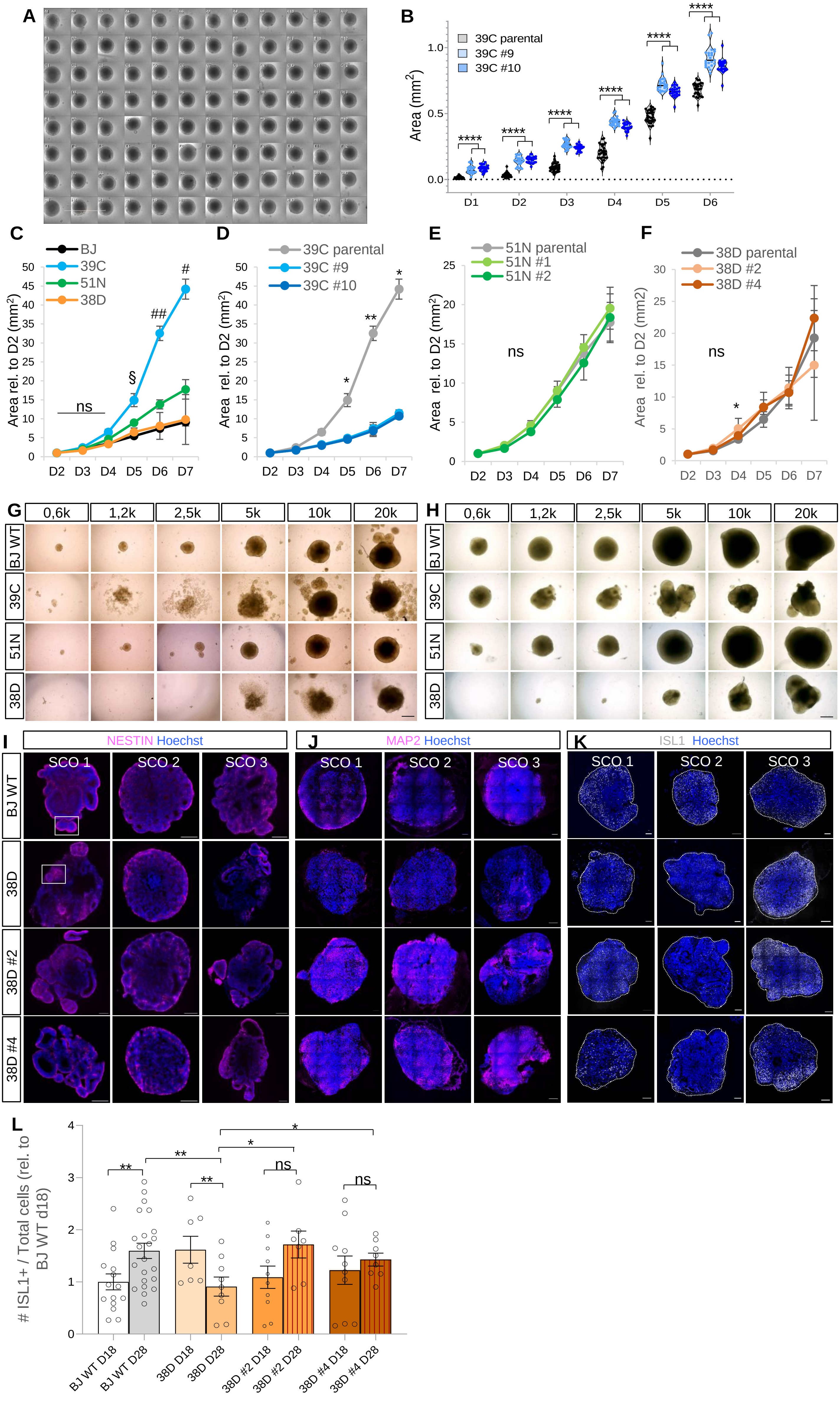

### Tables

| Primer type | Primer sequence |
|---|---|
| **Primers designed to detect non-targeted *SMN2*** | 1F GTTCTCCAAATCCGACCTCA |
| | 1R TTTCTTCCACATAACCAACCAG |
| **Primers designed to detect targeted SMN2** | 1F GCCCGGCCTAGTCTTGTATT |
| | 1R AGGTACCGTCGTCCTTGAAA |

**Table 1.** Primers used for PCR-amplification of SMN loci followed by Sanger sequencing.

| | Target Gene | Sequence forward (5´=>3´) | Sequence reverse (5´=>3´) |
|---|---|---|---|
| **hiPSC trilineage differentiation** | SOX1 | CTGACGTCCACTCTCAGTCT | CCACATCCTAATCTTGAGCCA |
| | PAX6 | TTGCCCGAGAAAGACTAGCA | TGGAGCCAGATGTGAAGGAG |
| | NCAM1 | GACCATCCACCTCAAAGTCTT | GAGGCTTCACAGGTAAGAGTG |
| | TBXT | CCACATAGTGAGAGTTGGGG | AGAGCTGTGATCTCCTCGT |
| | CXCR4 | AAATCTTCCTGCCCACCATC | GTACTTGTCCGTCATGCTTCT |
| | LEFTI1 | CTTGGGGACTATGGAGCTCAGG | ATGTACATCTCCTGGCGGC |
| | SOX17 | AACGCCGAGTTGAGCAA | GGCCGGTACTTGTAGTTGG |
| **SCO developmental gene expression** | SOX2 | GTACAACTCCATGACCAGCTC | CTTCAGCACCGAACCCAT |
| | NESTIN | CTCAGCTTTCAGGACCCCAAG | TCTCAAGGGTAGCAGGCAAG |
| | NGN2 | GCCAAAGTCACAGCAACG | TCCTCTTCCTCCTTCAACTCC |
| | DCX | GTGTTTATTGCCTGTGGTCCTG | GGAGGTTCCGTTTGCTGAGT |
| | SMI32 | GAGTGGTTCCGAGTGAGGCTG | AGTGAGTCCTTGGTGCTTTTCAG |
| | NKX6.1 | CCTGTACCCCTCATCAAGGA | GAATAGGCCAAACGAGCCCT |
| | HB9 | CTGGAGCACCAGTTCAAGCTCA | TGGAACCAAATCTTCACCTGGGT |
| | ISL1 | TGCTTTTCAGCAACTGGTCAAT | AGGACTGGCTACCATGCTGT |
| | CHAT | CGACAAGTCCCTGCATTTG | ACGGAGTCTGCTCGGATCA |
| **Housekeeping gene** | 18S | AAACGGCTACCACATCCAAG | CCTCCAATGGATCCTCCATA |

**Table 2.** Primers used for qPCR mRNA expression quantification for hiPSC Trillineage differentiation assay (genes recommended by STEMdiff Trilineage Differentiation Kit) and for neural differentiation of SCOs.
